# Proteolytic Control of an Auto-inhibitory Intrinsically Disordered Region Governs Small RNA Selectivity in Argonaute Proteins

**DOI:** 10.64898/2025.12.19.695459

**Authors:** Ida Isolehto, Lizaveta Pshanichnaya, Diego J. Páez-Moscoso, Maximilian Mager, Ann-Sophie Seistrup, Jan Schreier, Svenja Hellmann, Kumar Gaurav, Fridolin Kielisch, Juaxuan Chen, Lukas S. Stelzl, René F. Ketting

**Affiliations:** Institute of Molecular Biology, Mainz, Germany; PhD Programme on Gene Regulation, Epigenetics & Genome Stability, Mainz, Germany; Johannes Gutenberg University Mainz, Mainz, Germany; Max Perutz Labs, Vienna Biocenter Campus (VBC), Vienna, Austria; European Molecular Biology Laboratory, Epigenetics and Neurobiology Unit, Monterotondo, Italy

**Author notes:** Equally contributed (Ida Isolehto, Lizaveta Pshanichnaya, Diego J. Páez-Moscoso). Lead contact: RFK.

## Abstract

Argonaute proteins are central to small RNA-mediated gene regulation, yet the mechanisms controlling their activity remain incompletely understood. We elucidate a novel regulatory mechanism governing small RNA loading into the *C. elegans* Argonaute proteins WAGO-1 and WAGO-3. We show that N-terminal intrinsically disordered regions (N-IDRs) of these proteins do not affect subcellular localization but play critical roles in small RNA loading. We demonstrate that the N-IDR-processing protease DPF-3 facilitates small RNA loading in a catalysis-independent manner. Catalysis by DPF-3 and a second protease APP-1 is required, however, for activity. Deletion of these N-IDRs results in loading of aberrant small RNA species that trigger erroneous gene silencing. Supported by atomistic molecular dynamics simulations, we propose a model in which N-IDRs can simultaneously act as tuneable gatekeepers that auto-inhibit small RNA loading and as regulators of Argonaute stability, representing a previously unrecognized layer of regulation in Argonaute activity in small RNA pathways.

## INTRODUCTION

Argonaute proteins are deeply conserved proteins with important gene-regulatory functions^1^. These RNA binding proteins are guided to their target transcripts in a sequence-specific manner by their small RNA co-factors. They have been implicated in many different gene-regulatory processes, including the control of foreign nucleic acids in archaea^2^, virus control in plants^3^, transposon control in animals^4,5^ and developmental control of gene expression in both plants^6–8^ and animals^9,10^. Their relevance for human health became clear when mutations in AGO1 and AGO2, two Argonaute proteins, were identified in patients with neurodevelopmental disorders, now collectively known as Argonaute syndromes. Curiously, disease pathogenesis is primarily driven by gain-of-function mechanisms involving altered miRNA-mRNA targeting dynamics rather than loss of normal targeting function^11,12^. In addition, male infertility can be caused by mutations in Argonaute proteins or in processes closely linked to Argonaute protein activity^13,14^.

Metazoan Argonaute proteins are characterized by a specific set of folded domains. These are, from N- to C-terminus: N-domain, PAZ domain, MID domain and PIWI domain. Together, they fold into a bi-lobed structure, in which the 5’ end of the small RNA is bound by the MID domain and the 3’ end is bound by the PAZ domain^15^. The small RNA is presented in such a way that its 5’ region is pre-ordered for base-pairing, enhancing target site recognition. Following base-pairing between the small RNA and the target, the PIWI domain can be activated to cleave the targeted RNA molecule. However, not all Argonaute proteins contain cleavage-competent PIWI domains, and cleavage-independent target silencing through Argonaute proteins is a widespread phenomenon^5,16^.

On the N-terminal side of the N-domain, many Argonaute proteins contain an intrinsically disordered region (IDR). Such N-terminal IDRs (or N-IDRs) are poorly characterized despite their widespread prevalence, although a couple of examples have been described. In some Argonaute proteins, arginine methylation within the N-IDRs reflects their loading status^17^, and in other cases, N-IDRs contain nuclear localization signals^18,19^. However, the function of most N-IDRs remains unclear, and their importance may have been overlooked.

The nematode *Caenorhabditis elegans* (*C. elegans*) contains a large array of Argonaute proteins. Some can be recognized as clear homologs of mammalian Argonaute proteins, such as ALG-1 and ALG-2, which are functional homologs of the mammalian Ago1–4 proteins that act in the miRNA pathway^20^, or PRG-1, the single nematode homolog of the mammalian Piwi family^21–23^, which acts in the piRNA pathway. In addition to these well-conserved Argonautes, nematode genomes encode worm-specific Argonaute proteins, or WAGOs^16,24,25^. While one- to-one homologs cannot be identified in mammals, these WAGO proteins have all the structural features of other Argonaute proteins. For instance, 45% of Argonaute syndrome mutations in human AGO1/2 affect residues that are conserved in the WAGO protein WAGO-3, despite an overall sequence similarity of only ∼16%. At the functional level, WAGO proteins have specialized in driving the highly complex endogenous and exogenous RNA interference responses. Both the conserved structural features and the specialized roles of many WAGO proteins make them interesting targets for studies on molecular mechanisms of Argonaute proteins, as they allow manipulation of Argonaute functionality without, for instance, disrupting the developmentally essential miRNA pathway.

WAGO proteins bind small RNAs that are also known as 22G RNAs^26^. These small RNAs are made by RNA dependent RNA polymerases^27^, which are typically activated by another set of Argonaute proteins^28^. Hence, WAGO proteins and their bound 22G RNAs are also referred to as secondary Argonaute proteins and secondary small RNAs, respectively. Their individual functions at the molecular levels are not yet fully understood, but some are essential to trigger RNA interference (RNAi)^25,29^. This RNAi response can be inherited across generations in *C. elegans*^30–34^, and two particular WAGO proteins, WAGO-3 and WAGO-4, were shown to transmit the RNAi response from parent (male or female) to offspring^35,36^. For the paternal transmission through sperm, WAGO-3 requires the presence of the so-called PEI granules, formed by two spermatogenesis-specific proteins PEI-1 and PEI-2^37^. These PEI-granules physically ensure the presence of WAGO-3 in mature spermatids while excluding other Argonaute proteins during spermatid maturation.

Many of the WAGO proteins also contain N-IDR with unknown functions. Recent work showed that a dipeptidase of the DPPIV family, DPF-3, specifically, processes the N-IDR of WAGO-1 and WAGO-3^38^. Loss of DPF-3 activity had different effects on these two WAGO proteins. Whereas WAGO-1 started to bind 22G RNAs that are normally restricted to another WAGO-clade Argonaute (CSR-1), WAGO-3 was found to be destabilized in the absence of DPF-3. Also, transposon silencing was found to be crippled in *dpf-3* mutants. More recently, DPF-3 was implicated to act with ALG-1 and/or ALG-2, the two miRNA-binding Argonaute proteins^39^. In this study, DPF-3 was shown to bind ALG-1, and loss of DPF-3 activated ALG-2, whose N-terminal end was also shown to be a potential substrate of DPF-3. Curiously, these effects were independent of DPF-3’s catalytic activity. The mechanistic underpinnings of these effects of DPF-3 on Argonaute function have remained unclear, but the tight regulation of the N-IDRs suggests that these regions play a major regulatory role.

DPF-3 is a dipeptidyl peptidase, implying that it cleaves off di-peptides from the N-terminal end of polypeptides. An activity profile of DPF-3 was established using *in vitro* processing of a random set of polypeptides^40^. Its main activity was shown to be on N-termini with a proline at position two, but it can also act on dipeptides with other amino acids at position two, notably alanine. In contrast, the combined presence of prolines at positions two and three was shown to strongly inhibit its activity ^40^. These characteristics seem incompatible with the processing of the N-IDRs of WAGO-1 and WAGO-3, as DPF-3 would inevitably run into a position that would require it to cleave in between two prolines. Yet *in vivo*, WAGO-1 and WAGO-3 N-termini are processed much further^38^. Hence, the mechanisms underlying the processing of the WAGO N-IDRs and their regulatory roles are still unclear.

In this work we show that the N-terminal processing of WAGO-1 and WAGO-3 requires a second protease. This enzyme is named APP-1 in *C. elegans*, and it is the homolog of XPNPEP in humans. APP-1 is a peptidase that can remove an N-terminal amino acid when the second amino acid is a proline, and we previously identified it as an interactor of the RNAi-inheritance factor PID-2^41^. In our work, DPF-3 and APP-1 act successively on the N-termini of WAGO-1/3 to enable full processing, resembling the interplay between their human N-terminal protease homologs in inflammasome regulation^42,43^. DPF-3 also has a processing-independent function: it stimulates WAGO loading and is required for proper subcellular localization. Moreover, the N-IDRs themselves inhibit the unwanted loading of small RNAs onto WAGOs. Deleting the N-IDR of WAGO-1 and WAGO-3 causes binding of 22G RNAs that target histone genes. While these type of small RNAs are normally present at low levels, their elevated incorporation onto WAGO-1/3 triggers an RNAi cascade that leads to a reduction in histone mRNA levels, followed by severe defects in germline development. Structure predictions, followed by molecular dynamics simulations suggest that the N-IDR may modulate WAGO loading by physically interacting with the RNA-binding regions of the WAGO protein.

Overall, our data suggest a model for how the N-IDR of WAGO-1/3 dynamically controls Argonaute loading with small RNAs. We propose that the N-IDR may prevent the loading of small RNAs by blocking the RNA-binding surface of the Argonaute protein, until it is in an environment where DPF-3, and likely other factors, are present to enable Argonaute loading and activation. N-IDR-mediated Argonaute regulation has been described before in Archaea^44^, suggesting that this principle may be conserved among different domains of life.

## RESULTS

### DPF-3 alone does not fully process WAGO-1 and WAGO-3 N-IDRs

Previous work showed that the N-IDRs of WAGO-1 and WAGO-3 are processed *in vivo* by the dipeptidyl peptidase DPF-3^38^. This was correlated to the loading of WAGO-1 and stability of WAGO-3. However, only limited processing was observed *in vitro*, using a truncated version of recombinant DPF-3 and a synthetic N-IDR WAGO-1 peptide. We wanted to test if full-length recombinant DPF-3 (Figure S1A, lane 1) may process the N-IDRs further. However, incubation with WAGO-1 and WAGO-3 N-terminal peptides (assuming co-translational removal of the initiator methionine) still resulted in only restricted processing to amino acid positions 4 (His) and 6 (Val), respectively, even after 12 hours of incubation (Figure 1A, B top). The observed processing was specific to DPF-3 activity, as mutating one of the three residues of its catalytic triad (D861A; Figure S1A, lane 2) did not show any processing at any time point (Figure 1A, B bottom). We also noted that WAGO-3 processing was much less efficient than that of WAGO-1, consistent with the first dipeptide of WAGO-3 (Pro-Ala) being a sub-optimal DPF-3 substrate^40^. Indeed, when we incubated a WAGO-3 peptide starting at position 4 (Thr-Pro), we observed rapid removal of the Thr-Pro dipeptide (Figure S1B). However, processing then stopped again at Val6.

**Figure 1.**
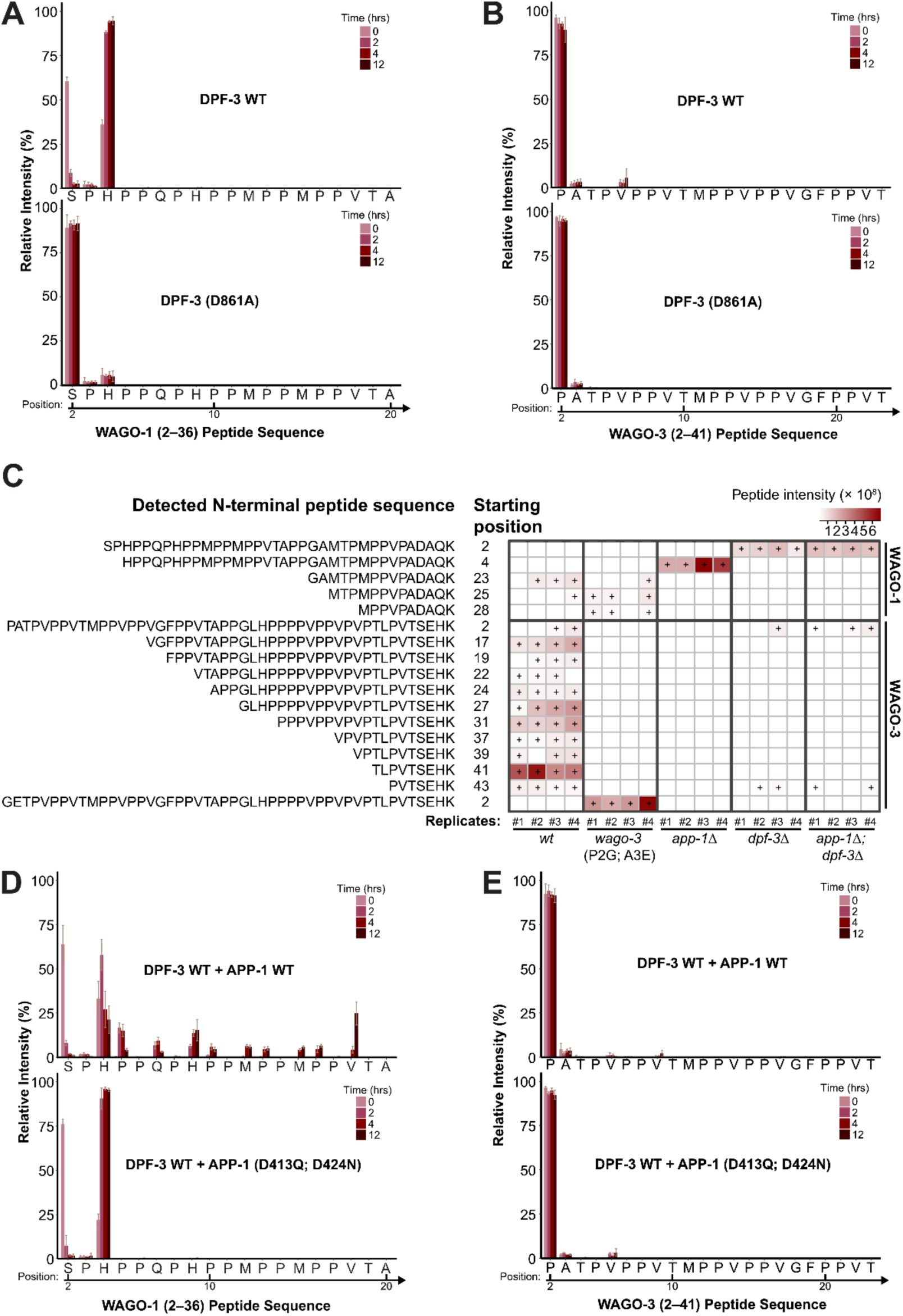
DPF-3 and APP-1 mediate N-terminal processing of WAGO proteins. (A and B) Bar plots showing mass spectrometry–based analysis of *in vitro* N-terminal cleavage of WAGO-1 (SPHPPQPHPPMPPMPPVTAPPGAMTPMPPVPADAQK) (A) and WAGO-3 (PATPVPPVTMPPVPPVGFPPVTAPPGLHPPPPVPPVPVPTL) (B) wild-type (WT) peptides incubated with recombinant WT DPF-3 (top) or the catalytically inactive DPF-3(D861A) mutant (bottom). Only a partial peptide sequence showing accumulation of processed fragments is shown on the X-axis. Numbers above the arrows denote the amino acid positions with respect to the full-length WAGO proteins. Time-course incubations (0, 2, 4, and 12 h) show that WT DPF-3 efficiently removes the first two amino acids of WAGO-1 peptides, whereas the DPF-3 mutant shows no detectable activity. In contrast, WT DPF-3 mediates limited cleavage, removing up to four residues, in WAGO-3 peptides. Bars represent means; error bars denote standard deviation (n = 2–3). (C) Heatmap summarizing the intensity of N-terminal peptides detected after Lys-C digestion from *in vivo* Argonaute protein enrichments in the indicated worm strains. Peptide identities and starting residue positions are shown on the left. The heatmap displays the LC-MS/MS intensity values for each peptide across individual replicates. A plus sign (+) indicates peptide detection despite low intensity. Loss of processing is observed in *wago-3(xe255)* (P2G, A3E), *dpf-3(xe68)*, and *app-1(xf323); dpf-3(xe68)* mutants. (D and E) As in (A) and (B), but with co-incubation of WAGO N-terminal peptides with WT DPF-3 and APP-1. The addition of WT APP-1 results in enhanced cleavage of WAGO-1 (D) and, to a lesser extent, WAGO-3 (E) peptides over time compared with reactions containing DPF-3 alone and catalytically inactive APP-1 (D413Q, D424N). This cooperative proteolysis depicts a functional interplay between DPF-3 and APP-1 in WAGO N-terminal processing. Bars represent means; error bars denote standard deviation (n = 2–3).

In contrast to these *in vitro* results, *in vivo* enrichment of Argonaute proteins followed by peptide profiling after Lys-C digestion, resulting in peptides terminating in lysine, revealed more extensive trimming of WAGO-1 and WAGO-3 N-termini in wild-type animals (Figure 1C). WAGO-1 showed a dominant N-terminal peptide starting at position 23 in three out of the four replicates, while the unprocessed peptide was not detected. For WAGO-3, unprocessed N-termini starting position 2 (in two replicates) and several processing intermediates were consistently observed, including a dominant peptide starting position 41 (Figure 1C). Many of these intermediate N-termini are predicted to be poor DPF-3 substrates^40^. In the absence of DPF-3, these WAGO-3 intermediate peptides were undetected (except for a low-abundance peptide starting at position 43), and for both WAGO-1 and WAGO-3 the N-terminal peptides that were detected corresponded to their unprocessed forms, confirming the role of DPF-3 in the processing of these N-IDRs (Figure 1C).

We conclude that DPF-3 processes the N-IDRs of WAGO-1 and WAGO-3, but its processing efficiency is not uniform along the N-IDRs. In fact, certain positions within these IDRs appear not to be processed by DPF-3.

### APP-1 processes Xaa-Pro-Pro sites to enable further DPF-3 processing

The discrepancies between *in vivo* and *in vitro* N-IDR processing suggested that additional factors might act in WAGO-1 and WAGO-3 N-IDR processing. The most significant block detected in our assay (Figure 1A, B; Figure S1B) occurred at N-termini starting with Xaa-Pro-Pro, consistent with the previously described DPF-3 activity profile^40^. In our previous work^41^ we identified APP-1, a highly conserved proline-specific peptidase (the nematode homolog of human XPNPEP1), as a protein interacting with the RNAi inheritance factor PID-2. APP-1 is expected to cleave in between any N-terminal amino acid and proline and could therefore potentially process intermediate N-terminal Xaa-Pro-Pro sequences where DPF-3 stalls. In fact, APP-1 has been shown to remove the N-terminal arginine from bradykinin (<u>**R**PP</u>GFSPFR)^45^. Such activity would then enable continued processing by DPF-3. To test this hypothesis, purified recombinant APP-1 (Figure S1A, lane 3) was incubated with a WAGO-1 peptide starting with His-Pro-Pro (mimicking a DPF-3 processing block; Figure 1A, top). Here, APP-1 efficiently processed the His-Pro-Pro N-terminus into a Pro-Pro N-terminus, which may then be further processed by DPF-3 (Figure S1C, top), while DPF-3 still could not process the His-Pro-Pro motif (Figure S1C, bottom). Note, however, that APP-1 slowly but appreciably also cleaved in between the two prolines after histidine removal from the His-Pro-Pro peptide. APP-1 alone could not process full length WAGO-1 or WAGO-3 N-terminal peptides *in vitro*, as no increased accumulation of shorter peptides was observed compared to the 0-hour time point (Figure S1D, E). For WAGO-1 this was unexpected, as the WAGO-1 N-terminus (Ser-Pro) is predicted to be an APP-1 substrate. Likewise, the WAGO-3 N-terminal peptide starting at position four (Thr-Pro) was not cleaved (Figure S1F), revealing unexpected substrate selectivity for APP-1. These results demonstrate differential and complementary substrate preferences for DPF-3 and APP-1. When incubated with both APP-1 and DPF-3 enzymes, WAGO-1 peptides were processed further than with either enzyme alone (Figure 1D, top). Additionally, processing intermediates of the WAGO-1 peptide clearly showed sequential cleavage activity between the two enzymes: while APP-1 removed one amino acid when the previous DPF-3 processing step resulted in a Xaa-Pro-Pro N-terminus, DPF-3 typically removed Pro-Pro dipeptides. The N-terminal WAGO-3 peptide was less efficiently processed (Figure 1E, top), likely due to the low DPF-3 processing efficiency of its N-terminus (Pro-Ala-Thr). Indeed, a synthetic peptide starting at position 4 was more efficiently processed, again revealing the above-mentioned alternate action between DPF-3 and APP-1 (Figure S1G, top). When key residues of the catalytic site of APP-1 were mutated (D413Q, D424N)^46^, the processing of both WAGO-1 and WAGO-3 peptides was similar to DPF-3-only processing (Figure 1D, E, bottom; Figure S1G, bottom). Finally, we note that both WAGO-1 and WAGO-3 N-IDRs contain sequences that are very poorly processed in vitro. Notably, these are Val-Thr, N-termini (Figure 1, S1). Because previous work has shown that these should be substrates of DPF-3^40^ we propose that these apparent *in vitro* processing blocks merely slow down the processing, but not stop it (also see Discussion).

To observe if APP-1 has the same role *in vivo*, WAGO-1 and WAGO-3 N-IDRs were profiled in *app-1* deletion animals. This showed a clear accumulation of WAGO-1 peptide starting at position 4 (His-Pro-Pro), consistent with the *in vitro* processing data (Figure 1C). On the other hand, no WAGO-3 peptides were detected at all (Figure 1C) due to *app-1* deletion affecting the steady-state levels of WAGO-3 (see below; Figure 4). Finally, absence of both APP-1 and DPF-3 phenocopied loss of only DPF-3 (Figure 1C). This is consistent with the first cleavage event being mediated by DPF-3 and suggests that the WAGO-3 intermediate that should accumulate in absence of APP-1 (starting at Val 6) is less stable than completely unprocessed WAGO-3. Taken together, our data strongly suggest that the combined action of APP-1 and DPF-3 generates the *in vivo* processing patterns of WAGO-1 and WAGO-3 N-termini.

### DPF-3 and APP-1 are required for RNAi inheritance

To analyse the roles of DPF-3 and APP-1 in RNAi, we first sequenced small RNA populations from both *app-1* mutants and *app-1;dpf-3* double mutants (Figure S2A,B) and also used previously published sRNA sequencing data for *dpf-3* mutants^38^. Analysis of overall 22G RNA compositions by sRNAseq revealed that loss of APP-1 and DPF-3 resulted in the loss of a large set of 22G RNAs. Upon further analysis, many of the lost 22G RNAs were described to be WAGO-1 and WAGO-3-associated^24^ (Figure S2C), consistent with their effects on these two Argonaute proteins, and consistent with conclusions drawn from the *dpf-3* single mutant data^38^. CSR-1-type 22G RNAs were largely unaffected (Figure S2D). The pools of affected 22G RNAs were very similar between the *app-1* deletion and catalytic mutants, especially those that were lost (Figure S2E). Between *app-1* and *dpf-3* mutants the results were also similar, although here more unique 22G RNA targets were identified (Figure S2F).

We originally discovered APP-1 in a study addressing transposon silencing and RNAi inheritance^41^. We showed it interacts with PID-2, a protein involved in RNAi inheritance and transposon silencing, but a role in these processes for APP-1 itself remained unclear. Likewise, while DPF-3 has been implicated in transposon silencing^38^, its role in RNAi inheritance is unclear. We found that both factors are required for the silencing of Tc1 excision in the germline (Figure 2A). Furthermore, we tested *app-1* and *dpf-3* deletions for RNAi inheritance using two different assays. First, we fed hermaphrodites expressing GFP in the oocytes with food that triggers RNAi against GFP and scored the silencing in both the fed animals (P0) as well as their F1 progeny which were not on RNAi any longer, using *hrde-1* as a control, being a known RNAi-inheritance factor^34^. This revealed a clear RNAi inheritance defect (Figure 2B). Second, we used a recently developed, parent-specific RNAi inheritance scheme^35^. We triggered RNAi against GFP in a homozygous mutant male, which was subsequently crossed to naive hermaphrodites, followed by raising the F1 to adulthood in absence of dsRNA triggers. GFP expression was measured in both P0 and F1 to assess RNAi inheritance. This experiment revealed a clear loss of RNAi inheritance from both *app-1* and *dpf-3* mutant males in F1 animals (Figure 2C–E). Moreover, catalytically inactive *app-1* and *dpf-3* mutants, as well as a non-processive *wago-3*(P2G; A3E) variant, showed similar inheritance defects. We conclude that both APP-1 and DPF-3 enable RNAi inheritance through both sperm and oocyte.

**Figure 2.**
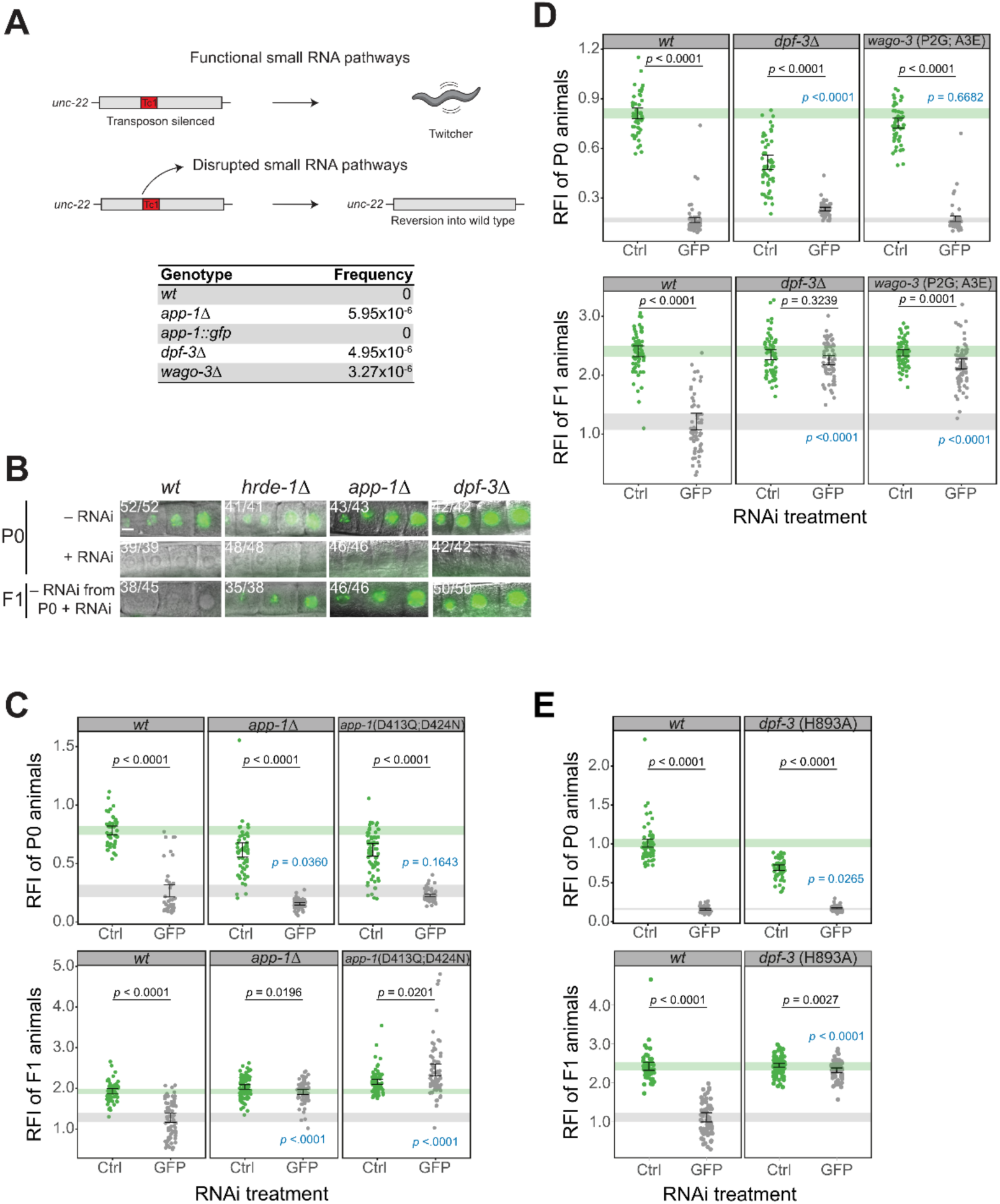
N-terminal processing of WAGO-3 is required for RNAi inheritance. (A) Tc1 transposon excision frequency in *app-1*, *dpf-3* and *wago-3* mutants. *unc-22(st136)* animals carrying Tc1 insertions were grown until starvation. This insertion causes a clearly visible locomotion defect. Excision of the Tc1 element in the germ cells can restore *unc-22* function in the offspring, leading to normal locomotion. Animals with such restored locomotion were quantified, as a measure of Tc1 excision frequency, being a proxy of Tc1 activity. (B) Inheritance of *gfp* RNAi-induced silencing. Representative fluorescence images of P0 and F1 animals expressing GFP::H2B in germline nuclei. P0 animals were exposed to *gfp* RNAi (RNAi+) or empty vector control (RNAi-). F1 progeny were cultured and scored on OP50 plates. Numbers indicate animals with the depicted phenotype out of total scored. Scale bar, 10 μm. (C) - (E) Paternal inheritance of *gfp* RNAi-induced silencing. RNAi-treated P0 males were crossed to untreated hermaphrodites. Male F1 cross-progeny were scored for GFP silencing. Relative fluorescence intensity (RFI) was calculated as GFP signal normalized to RFP signal for each animal. n = at least 45 animals per condition. Error bars show 95% confidence intervals for the mean. Shading indicates the 95% confidence interval of untreated (green) and *gfp* RNAi treated (grey) wild-type for comparison across genotypes. Statistical significance was assessed testing contrasts in Gaussian linear models. Black *p*-values: effect of RNAi treatment. Blue *p*-values: genotype effect in GFP RNAi treated group. *P*-values corrected for multiple testing within each contrast type.

### DPF-3 and APP-1 affect WAGO-3 localization in PEI granules

We recently identified inheritance-related germ granules (PEI granules) that form during spermatogenesis^37^. These contain WAGO-3, which is essential for RNAi inheritance via sperm^35^. However, the initial studies on PEI granules used N-terminally tagged WAGO-3, preventing WAGO-3 processing. Thus, we first tested how internally tagged WAGO-3 behaved. In general, WAGO-3 was less abundant when tagged internally versus N-terminally (Figure S3A), but the tagging site did not appear to strongly affect subcellular localization patterns: WAGO-3 was still observed in perinuclear granules in the gonads (Figure S3C) and was still enriched within the PEI granules that form during spermatogenesis (Figure 3A). We noted, however, that there was relatively strong diffuse WAGO-3 signal in the cytoplasm of the gonad (Figure S3C,D) and in the residual body during spermatogenesis (Figure 3A) when internally tagged, possibly related to its lower expression level in general. We next tested if APP-1 and DPF-3 affected the localization of WAGO-3 in spermatids and found that absence of either protein resulted in alterations of WAGO-3 localization (Figure 3A). In *dpf-3* deletion mutants, WAGO-3 signal was weak and diffuse, whereas upon *app-1* deletion WAGO-3 was found in cloud-like structures within the spermatids (Figure 3A). These were still dependent on PEI-1, as they disappeared upon *pei-1* deletion (Figure 3A). These results suggest that WAGO-3 associates less stably with the PEI granules in the absence of APP-1. In *app-1;dpf-3* double mutants, WAGO-3 signal also appeared mostly diffuse throughout the spermatids and residual body (Figure 3A).

**Figure 3.**
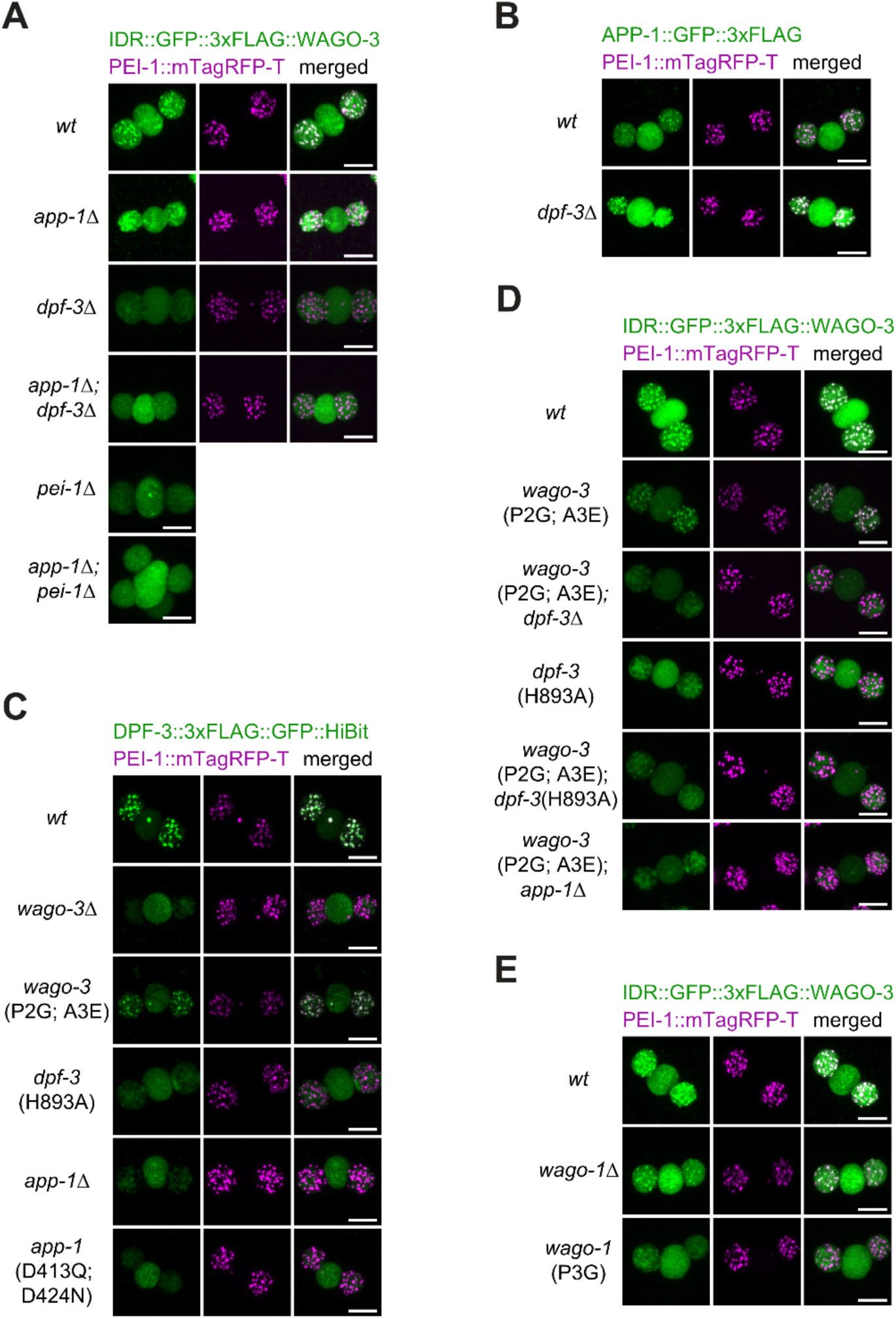
Localization of WAGO-3 to PEI granules depends on DPF-3 and APP-1. A), D), and E) Representative confocal images of dissected *C. elegans* budding spermatids showing IDR::GFP::3xFLAG::WAGO-3 (green), and PEI-1::mTagRFP-T (magenta) localization in the specified mutants. All strains additionally carry the *him-5(e1490)* mutation in the background. Maximum intensity projections. Scale bar, 5 μm. Images are representative of at least 10 budding spermatids pooled from at least two independent experiments. B) As above except for APP-1::GFP::3xFLAG (green), C) As above except for DPF-3::3xFLAG::GFP::HiBit (green).

To determine the localization of APP-1, GFP was inserted at its C-terminus. The tagging did not strongly affect APP-1 function, as silencing of Tc1 excision in the germline was comparable to that in the wild type (Figure 2A). Interestingly, both APP-1 and DPF-3 themselves are enriched in PEI granules (Figure 3B, C). While APP-1 localization to the PEI granules did not depend on DPF-3 (Figure 3B), DPF-3 required both APP-1 and WAGO-3 (Figure 3C; APP-1 localization in absence of WAGO-3 was not tested). To assess if WAGO-3 processing was required for this, we tested whether the confirmed non-processable WAGO-3 (P2G;A3E) variant (see Figure 1C) could still support DPF-3 enrichment in PEI granules and found that it supported normal DPF-3 localization (Figure 3C). Reversely, WAGO-3(P2G;A3E) also required DPF-3 for proper PEI granule localization, indicating that this mutual effect between WAGO-3 and DPF-3 is independent of DPF-3 processing the WAGO-3 N-IDR (Figure 3D). However, DPF-3 did not enrich in PEI granules when catalytically inactive itself (Figure 3C), and WAGO-3(P2G;A3E) localization was affected by a catalytic mutation in DPF-3, just like wild-type WAGO-3 (Figure 3D, Figure S3B). Our interpretation of these seemingly contradictory results is that catalytically inactive DPF-3 may become trapped on other substrates and therefore less available to enable proper WAGO-3 localization. As WAGO-1 is another DPF-3 substrate, we reasoned that non-processable WAGO-1 may thus interfere with WAGO-3 localization, as it may titrate DPF-3 away from WAGO-3. Indeed, while deletion of *wago-1* did not affect WAGO-3 presence in PEI granules, a processing-defective version of WAGO-1 (WAGO-1(P3G))^38^ resulted in absence of WAGO-3 from the PEI granules (Figure 3E). These results depict a competitive scenario between DPF-3 substrates, and interactions between DPF-3 and its substrates that are more stable than mere enzyme-substrate interactions.

To identify stable DPF-3 interactions, we performed DPF-3 immuno-precipitation (IP) from males followed by mass spectrometry. Consistent with previous work^39^, the Argonaute protein ALG-1 was enriched in DPF-3 IPs (Figure S4), supporting the idea of stable DPF-3-Argonaute interactions. In contrast, however, WAGO-1 and WAGO-3 were not enriched (Figure S4), indicating that WAGO-1 and WAGO-3 do not interact with DPF-3, or at least not as stably as ALG-1. We conclude that WAGO-1 and WAGO-3 interact with DPF-3 in a processing-independent manner, but that these interactions are not stable, or abundant enough to be detected in immuno-precipitation experiments.

### DPF-3 and APP-1 affect WAGO-3 stability and loading in a processing-independent manner

DPF-3 was shown before to stabilize WAGO-3^38^. We addressed this aspect again, now also including *app-1* mutants. In addition, we focussed our analysis on males to allow comparison with the above-described subcellular localization during the final stages of spermatogenesis. On Western blots, both *dpf-3* and *app-1* deletions reduced WAGO-3 abundance (Figure 4A, lane 2 and 3; B, lane 2; D), consistent with previous findings in hermaphrodites. Interestingly, WAGO-3(P2G;A3E) abundance was also affected by DPF-3 (Figure 4B, lane 3 and 4;C, lane 4 and 5; D), showing that the decrease in protein levels was not (only) related to the processing of the N-IDR, but also depends on the presence of DPF-3, independently of its processing activity.

**Figure 4.**
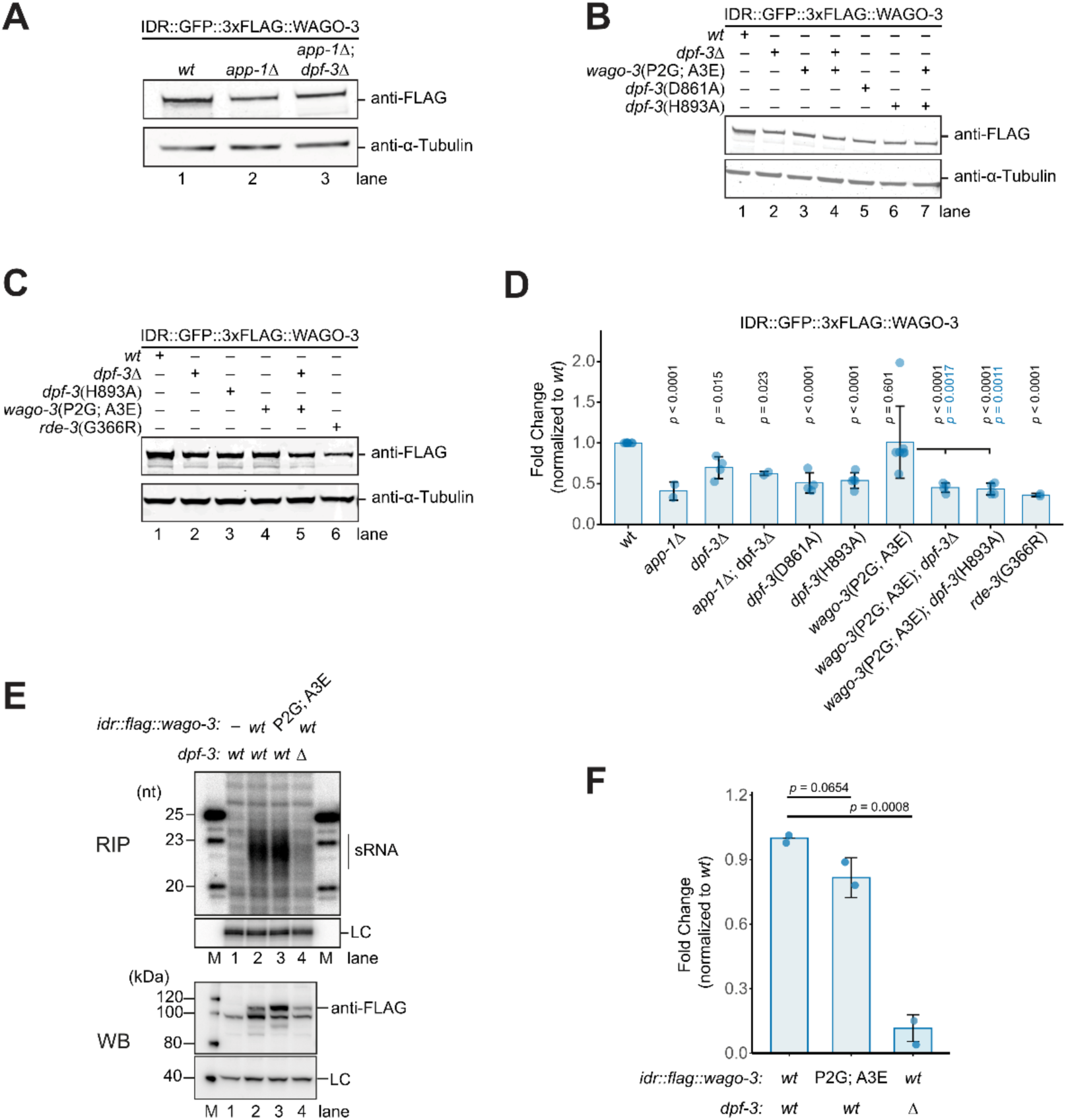
WAGO-3 stability and loading are influenced by APP-1 and DPF-3 independently of WAGO-3 processing. (A) Endogenous WAGO-3 protein levels in *app-1* and *dpf-3* double deletions. Representative Western blot comparing levels of internal GFP::3xFLAG-tagged WAGO-3. Denatured whole-worm protein extracts obtained from adult males carrying the *him-5(e1490)* mutation and PEI-1::mTagRFP-t in the background were resolved by SDS-PAGE. The blot was probed with anti-FLAG to detect WAGO-3 (top) and with anti–α-tubulin (bottom) as a loading control. Relative quantification of WAGO-3 is shown in (D); n = 2. (B) Endogenous WAGO-3 protein levels in *dpf-3* mutants and/or non-processive wago-3 mutants. Representative Western blot comparing levels of internal GFP::3xFLAG-tagged WAGO-3. Protein levels were assessed as in (A). Relative quantification of WAGO-3 is shown in (D); n = 2–4. (C) Endogenous WAGO-3 protein levels in *dpf-3* mutants and/or non-processive wago-3 mutants together with *rde-3* mutant. Representative Western blot comparing levels of internal GFP::3xFLAG-tagged WAGO-3. Protein levels were assessed as in (A). Relative quantification of WAGO-3 is shown in (D); n = 2. (D) Quantification of the effect of *app-1, dpf-3, wago-3,* and *rde-3* mutants on WAGO-3 protein levels. Bar plot showing fold-change values relative to endogenously IDR::GFP::3xFLAG-tagged WAGO-3. Quantifications are based on the blots shown in panels A–C. Black *p*-values indicate statistical significance of effects relative to *wt* using empirical Bayes-moderated one-sample t-tests implemented in Limma; corrected for multiple testing using Holm’s method. Blue *p*-values correspond to fold change comparisons to *wago-3*(P2G; A3E), tests of contrasts in a linear model, corrected for multiple comparisons against one control (multivariate t-method). Bars represent mean ± SD (n = 2–4). (E) RNA loading assay by RNA immunoprecipitation (RIP) from FLAG-tagged WAGO-3 in adult males. To detect background levels, a sample containing untagged WAGO-3 was included in the assay (lane 1). (Top) RNA was isolated, dephosphorylated with CIP, 5′-end– radiolabeled with [γ-³²P]ATP using T4 PNK, and analyzed on a 12% urea, 8 M denaturing PAGE gel. A [γ-³²P]ATP–labeled 86-mer oligonucleotide was used as a loading control (LC), and a size ladder was run on both edges of the gel; marker sizes are indicated on the left. (Bottom) Levels of soluble WAGO-3 from the input samples shown above were analyzed by western blot using an anti-FLAG antibody. A non-specific band recognized by the anti-FLAG antibody in the absence of the epitope tag (lane 1) served as an intrinsic loading control. (F) Bar plot showing the relative quantification of RNA immunoprecipitated with WAGO-3 in different genetic backgrounds, based on the blots shown in (E). RNA fold enrichment is presented relative to FLAG-tagged WAGO-3 after normalization to the protein levels in the input. Black *p*-values indicate statistical significance as described in (D). Bars represent mean ± SD (n =2).

One known aspect that can affect Argonaute stability is its loading with small RNAs. Indeed, WAGO-3 levels dropped in *rde-3* mutants, in which no 22G RNA co-factors for WAGO proteins are made (Figure 4C, lane 6; D). Based on these results we hypothesized that WAGO-3 is unstable in *dpf-3* mutants because it fails to be loaded. To test this, we immunoprecipitated internally tagged WAGO-3 from wild-type and *dpf-3* deletion backgrounds, and probed WAGO-3 loading by analysing the total bound sRNA pool on a sequencing gel (Figure 4E, top). When we normalized the detected sRNA amounts to the amount of WAGO-3 protein present in the native protein extract, we found that loss of DPF-3 triggered a severe WAGO-3 loading defect (Figure 4E,F). Meanwhile, the non-processable WAGO-3(P2G;A3E) was loaded at close to wild-type levels (Figure 4E, lane 3). This shows that DPF-3 affects the loading of WAGO-3 in a processing-independent manner and suggests that the reduced stability of WAGO-3 in absence of DPF-3 is related to ineffective loading of WAGO-3 with small RNAs.

### N-IDR processing is required for RNAi inheritance

What then is the role of the processing of the IDR? We focused on WAGO-3 for this question because of its well-defined function in RNAi inheritance and asked whether males that express WAGO-3(P2G;A3E) have a defect in RNAi inheritance, despite the fact that this protein can be loaded with small RNAs and is found in the PEI granules. Strikingly, males expressing WAGO-3(P2G;A3E) were fully defective in RNAi inheritance (Figure 2D), showing that the processing of the N-IDR of WAGO-3 is likely required at a stage following its localization in PEI granules during sperm maturation. Possibly this relates to WAGO-3 protein stability being affected by an unprocessed N-IDR (see Discussion and Deng *et al*.(parallel submission)).

### Deletion of N-IDRs from WAGO-1 and WAGO-3 triggers sterility

Given the above results, we next asked if deletion of the complete N-IDRs affected WAGO loading with 22G RNAs. Initial genome editing experiments on the *wago-3* N-IDR indicated acute dominant sterility, preventing the establishment of a strain with this deletion. In contrast, full deletion of *wago-3* does not display such an acute phenotype^37^. Because of this result, we deleted the N-IDR regions from WAGO-1 and WAGO-3 (WAGO-1/3(ΔIDR)) in a setting where we could repress and release their expression in a controllable manner. To achieve these two aims, we tagged wild-type and N-IDR deletion (ΔIDR) versions of WAGO-1 (deletion of the first 37 amino acids) and WAGO-3 (deletion of the first 50 amino acids) with auxin-inducible degrons, allowing us to culture animals in presence of auxin and to induce their expression by auxin withdrawal (Figure 5A-B)^47^. WAGO-1 was tagged internally, and WAGO-3 was tagged N-terminally, but both approaches yielded similar results. When grown on auxin-containing media, WAGO-1/3(full-length) and WAGO-1/3(ΔIDR) strains were viable and fertile. However, when L1-stage larvae were put onto plates lacking auxin, WAGO-1/3(ΔIDR) animals developed into sterile adults, consistent with our initial genome editing efforts. When analysed in more detail, we noticed that WAGO-1/3(ΔIDR)-expressing animals contained underdeveloped germlines (Figure 5C-D). Whereas germline formation still appeared normal at the L2 stage, examination at the L3 stage demonstrated that expression of WAGO-1/3(ΔIDR) resulted in aberrant germline development (Figure S5A-B).

**Figure 5.**
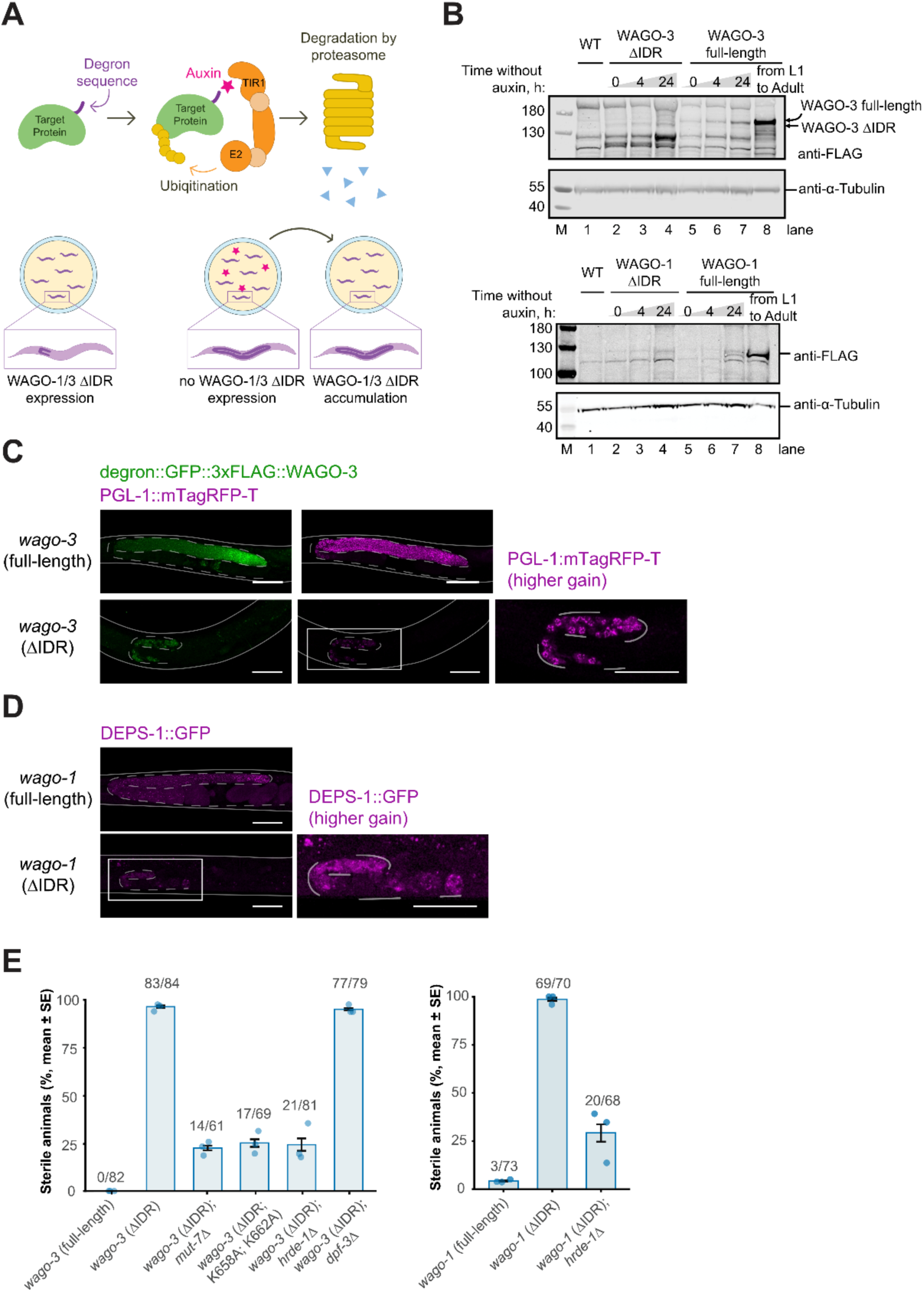
Deletion of N-IDRs from WAGO-1 and WAGO-3 triggers sterility. (A) Schematic representation of auxin-inducible protein degradation system. Expression of WAGO-3(ΔIDR) or WAGO-1(ΔIDR) on auxin-free plates results in germline sterility. To enable experimental analysis, animals were maintained on auxin-supplemented plates until the adult stage, then transferred to auxin-free plates to permit accumulation of ΔIDR variants. (B) Immunoblot analysis of FLAG-tagged WAGO-3 or WAGO-1 accumulation kinetics following auxin removal. Whole-worm protein extracts from gravid adult hermaphrodites were resolved by SDS-PAGE and probed with anti-FLAG antibody. α-tubulin served as a loading control. n = 2. (C-D) Representative fluorescence microscopy of degron::GFP::3×FLAG::WAGO-3 and PGL-1::mTagRFP-T (C) and DEPS-1::GFP (D) in the indicated mutants. Solid lines represent the worm body, and dashed lines represent the germline boundary. The white box indicates a magnified region. Images are representative of three independent biological replicates. Scale bars, 50 μm. (E) Quantification of sterility in animals cultured on auxin-free plates. Bars indicate the percentage of sterile hermaphrodites; numbers above bars denote sterile animals/total animals examined for each strain. Error bars represent the standard error of the mean.

The strong phenotype of WAGO-3(ΔIDR) animals could be fully rescued by combining it with a *mut-7* mutation, which strongly reduces 22G RNA biogenesis^48,49^ (Figure 5E, Figure S5D), suggesting that the phenotype of IDR deletion is driven by erroneously bound 22G RNAs. To test this further, we also aimed to examine an effect of WAGO-3 RNA-binding defective mutant. To design such a mutation, we ran atomistic molecular dynamics simulations (8 x 1.5 µs) (Table S1) starting from an AlphaFold3-predicted structure of the WAGO-3 RNA complex. These simulations showed many contacts between the RNA and the central cavity (Figure S5C). For instance, Ala 323, Arg 423, Ile 614 and Lys 877 showed high contact probability with the RNA. Within the MID domain, Lys 658 showed strong contact. A close-by residue, Lys 662, is additionally likely involved in 5’ phosphate coordination based on structural information from another Argonaute protein^50^. We decided to jointly mutate the Lys658 and Lys662 residues to alanine to interfere with 22G RNA binding. An indication that WAGO-3(K658A;K662A) indeed did not bind RNA is that it was less stable than wild-type WAGO-3 (Figure S5F). Furthermore, it accumulated in Mutator foci (see below; Figure 6E), where MUT-7 is present to help 22G RNA biogenesis^51,52^. Consistent with the result for *mut-7* mutation, WAGO-3 RNA-binding mutation fully rescued the ΔIDR phenotype (Figure 5E, Figure S5D).

**Figure 6.**
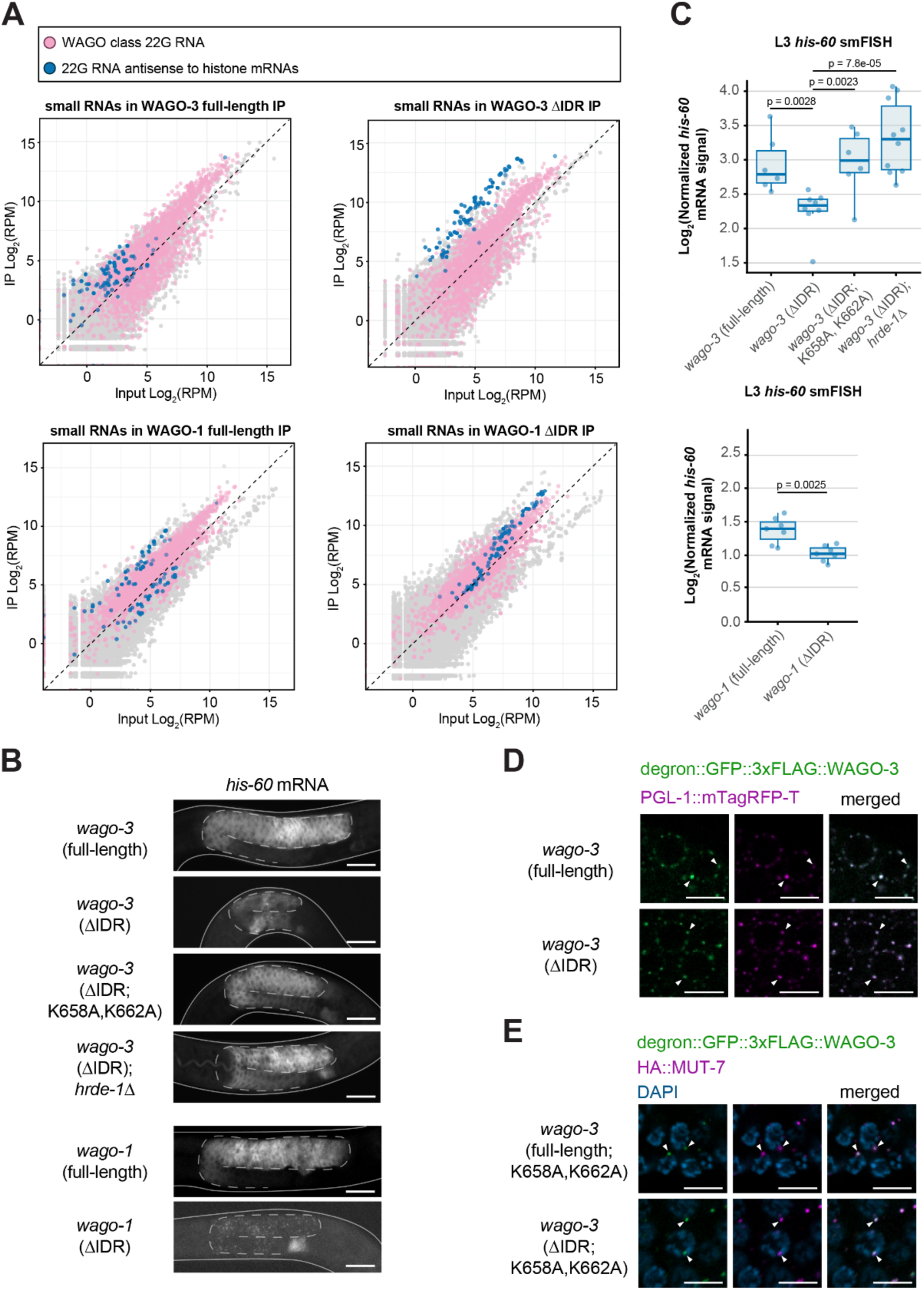
N-IDR deletion triggers loading of histone-targeting 22G RNAs into WAGO-1 and WAGO-3. (A) Scatter plots depicting 22G-RNA enrichment in WAGO-3 or WAGO-1 immunoprecipitation. Log₂-transformed reads per million (RPM) values for 22G-RNAs mapping to individual genes in IP samples (y-axis) are plotted against input samples (x-axis) from day 1 adults following 4 hours of auxin withdrawal. Canonical WAGO target genes and histone genes are indicated in pink and blue, respectively. n = 4 biological replicates per genotype. (B) Maximum intensity projections of confocal *his-60* smFISH in L3-stage larvae cultured on auxin-free plates. Solid lines outline the worm body and dashed lines outline the germline. Images are representative of six biological replicates. Scale bar, 20 μm. (C) Quantification of normalized his-60 mRNA smFISH signals in L3-stage larvae shown in (B). Each dot represents an individual animal; horizontal bars indicate mean values, and error bars represent standard deviation. Statistical significance was assessed using Welch’s t-test for WAGO-1 comparisons and Dunnett’s multiple comparison test for WAGO-3 comparisons. n = 6 biological replicates per genotype. (D) Representative fluorescence micrographs of the distal germline region in day 1 adult hermaphrodites expressing degron::GFP::3×FLAG::WAGO-3 and PGL-1::mTagRFP-T in the indicated genetic backgrounds, imaged 24 hours after auxin removal. Arrows indicate individual perinuclear germ granules. Scale bars, 5 μm. (E) Immunofluorescence imaging of the distal germline region from dissected day 1 adult gonads expressing degron::GFP::3×FLAG::WAGO-3 (K658A;K662A) and HA::MUT-7 in the indicated genetic backgrounds. Samples were probed with anti-GFP and anti-HA antibodies. Arrows indicate individual perinuclear germ granules. Scale bars, 5 μm.

To further probe the idea that erroneous endogenous RNAi drives the ΔIDR phenotype, we also tested whether loss of HRDE-1 could rescue the phenotype triggered by N-IDR deletion, since our recent work suggests that HRDE-1 can work downstream of WAGO-3 in establishing silencing^35^. Indeed, *wago-1(*Δ*IDR);hrde-1* and *wago-3(*Δ*IDR);hrde-1* double mutants were fertile, indicating that off-targeting of the endogenous RNAi machinery drives the sterility phenotype, and not an RNAi-unrelated toxic effect of WAGO-1/3(ΔIDR) proteins (Figure 5E, Figure S5D). Finally, given the role of DPF-3 in WAGO-3 loading we also tested *wago-3(*Δ*IDR);dpf-3* double mutants. These were still sterile, and in fact showed an even more severe germline developmental defect (Figure 5E, Figure S5D). This suggests that when WAGO-3 does not have an N-IDR, DPF-3 is not required for WAGO-3 loading.

We conclude that the N-IDRs can block the loading of erroneous small RNAs into WAGO-1 and WAGO-3 and suggest that DPF-3 plays a role in relieving this block in surroundings where the ‘proper’ WAGO-1/3 22G RNAs are being made.

### N-IDR deletion triggers loading of histone-targeting 22G RNAs into WAGO-1 and WAGO-3

Next, we aimed to sequence the 22G RNAs bound to WAGO-1 and WAGO-3, with or without their N-IDRs present. However, given the strong developmental phenotypes, releasing WAGO-1/3(ΔIDR) expression from larval stages onwards would lead to convoluted results. Hence, to minimize chances of secondary effects, we let hermaphrodites develop until adulthood on auxin and then transferred them to non-auxin media for four hours (Figure 5B), before harvesting the animals for immunoprecipitation. Both proteins bound their previously described^24^ 22G RNA populations, irrespective of the presence of their N-IDR (Figure 6A). However, in addition to these ‘regular’ 22G RNAs, we detected a significant increase in 22G RNAs that target replication-dependent histone transcripts. This increase was also detectable in the input RNA, suggesting that expression of WAGO-1(ΔIDR) or WAGO-3(ΔIDR) leads to the stabilization of such 22G RNA species within these mutated WAGO proteins (Figure S6A). Also, given the fact that HRDE-1 is required to drive the phenotype, a part of these histone-targeting 22G RNAs will likely be bound to HRDE-1, contributing to their elevated levels overall.

The under-developed germline phenotype is consistent with the silencing of histone mRNAs at a stage when the germline proliferates. To probe this, we performed smFISH against *his-60* mRNA. This revealed that, indeed, histone mRNA levels dropped at the L3 stage, when the phenotype developed (Figure 6B-C, Figure S6B-C). We conclude that the N-IDRs of WAGO-1 and WAGO-3 prevent the loading of histone-targeting 22G RNAs and the consequent erroneous silencing of histone expression during germline development.

### N-IDR deletion of WAGO-3 does not affect its sub-cellular localization

IDRs of proteins are often linked to phase-separation-related processes^53–55^. Thus, WAGO-1 and WAGO-3 could be loaded with erroneous small RNAs because their sub-cellular localization may be affected by the absence of their N-IDRs. We therefore probed the effect of the N-IDR on subcellular localization. Following the logic described above for RNA immunoprecipitation experiments, we induced WAGO expression for twenty-four hours in adult hermaphrodites before analysis (Figure 5B), focusing on WAGO-3. These analyses revealed that WAGO-3 localization remained largely unaffected by deletion of its N-IDR (Figure 6D-E, Figure S6E). WAGO-3 was detected in P-granules throughout the germline, as demonstrated by its co-localization with PGL-1. Moreover, when we abolished WAGO-3 loading by mutation of residues in its MID domain, no differences were observed between WAGO-3 with or without its N-IDR: in both cases, the protein colocalized with MUT-7, which served as a Mutator foci marker. WAGO-3 co-localization with PEI-1 in budding spermatids were also not affected by IDR deletion (Figure S6D). We conclude that the steady-state subcellular localization of loaded and unloaded WAGO-3 is not determined or affected by its N-IDR.

### The N-IDR occupies the RNA-binding groove in molecular dynamics simulations

How can the N-IDR prevent the loading of histone-targeting 22G RNAs? One option that we considered is that the N-IDR may prevent the loading of any small RNA until the WAGO protein is in an environment in which such an auto-inhibitory function can be counteracted. Potentially, such a mechanism could provide a tight coordination between 22G RNA production and WAGO loading in time and space.

When we analysed WAGO-1 and WAGO-3 structures on AlphaFold2^56^ we noticed two things. First, the end of the N-IDR is characterized by an alpha-helix (helix^1^) in WAGO-1 and WAGO-3, and this helix docks onto the body of the Argonaute fold, close to the beginning of the RNA-binding regions (Figure S7A). Such a helix^1^ was also found on the closely related WAGO-4 protein but not on any other Argonaute proteins in *C. elegans* we looked at (Figure S7A). Helix^1^ brought the N-IDR close to the RNA-binding region of WAGO-1 and WAGO-3. Second, we noted that the N-IDR was often placed within the RNA-binding groove WAGO-3 when no 22G RNA was provided (Figure 7A). In this conformation it was often close to another alpha-helix: helix^2^. When a 22G RNA sequence was provided the N-IDR was placed outside this groove (Figure 7B). To probe the stability of the predicted conformation with the N-IDR within the RNA-binding groove, we further analysed the WAGO-3 predictions using atomistic molecular dynamics simulations (Table S1). Firstly, this revealed that Helix^1^ is stably maintained over a period of 5 µs, as seen in four simulations started from two different predicted structures (Figure S7B,C; Movie S1). Secondly, the N-IDR in fact docked stably into the RNA-binding groove (Figure 7C). In these four simulations the distance between N-IDR Residue 34 to helix^2^ (Figure 7A) fluctuated between ∼10 Å and ∼20 Å (Figure 7C; Movie S1). In contrast, N-IDRs starting outside this RNA-binding groove fluctuated at a distance 30 Å and 70 Å (Figure 7C). Interestingly, many residues that we defined before to interact with RNA, also interacted with the N-IDR (Lys 321, Arg 441, Arg 443, Lys 457, Arg 466, Arg 809) (Figure 7D). Notably, also Pro-Gln interactions were observed, in particular between the Pro residues 30-33 within the N-IDR of WAGO-3 and the Gln residues 453 and 873 on the surface of the RNA-binding groove (Figure S7D-E).

**Figure 7.**
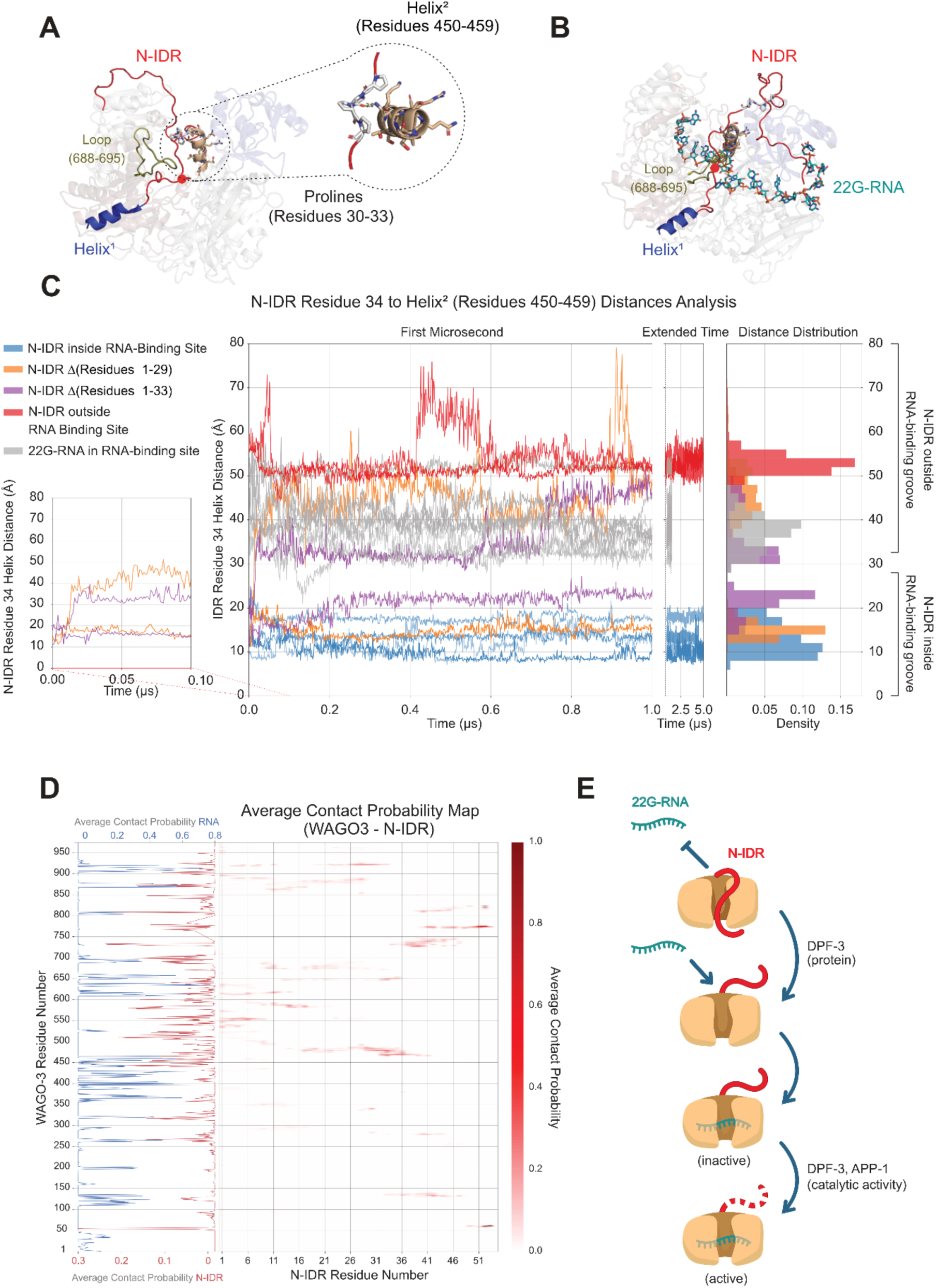
Structural dynamics of WAGO-3 IDR / 22G-RNA interactions analyzed by molecular dynamics simulations. (A) Representative snapshot of a molecular dynamics (MD) simulation of wild-type (wt) WAGO-3 where the N-IDR is positioned inside the 22G-RNA binding cleft. The interacting helix^2^ (residues 450-459) is represented in wheat. The inset shows a close-up of the N-IDR proline cluster engaging glutamine residue 453 within the helix^2^. The N-IDR is shown in red, with helix^1^ highlighted in blue, the interacting helix^2^ (residues 450-459) is represented in wheat, loop (residues 688-695) is in green, the red point represents the end of N-IDR processing. (B) Snapshot of a wt WAGO-3 MD simulation with a 22G-RNA bound in the canonical binding cleft. In this state, the N-IDR (red) is positioned outside the RNA-binding site. Helix^1^ is highlighted in blue, the interacting helix^2^ (residues 450-459) is represented in wheat, loop (residues 688-695) is in green, the red point represents the end of N-IDR processing. (C) Multi-group comparison of distances between N-IDR residue 34 and helix^2^ residues 450-459 across simulations lasting 1-5 μs. Different simulation conditions are color-coded, individual replicates are indicated by transparency. The zoom in on the first 100 ns highlights simulations in which the first 29 or 33 residues of the N-IDR were truncated and a shift in N-IDR positioning compared to the wt was visible. (D) Contact probability maps averaged over four wt WAGO-3 simulations, showing residue-level contact probabilities between WAGO-3 and the N-IDR (heatmap color coded from 0 (no contact) to 1.0 (persistent contact)). The accompanying line plot compares contact probabilities of WAGO-3 residues with the N-IDR and with 22G-RNA of eight simulations. (E) Schematic representation summarizing steps of WAGO-1/3 loading with 22G-RNA. DPF-3 and WAGO protein-protein interactions promote WAGO loading. DPF-3 catalytic function is necessary for WAGO-1/3 N-IDR processing and RNAi inheritance.

Perhaps not surprisingly, the RNA-interactions were significantly more persistent, suggesting that RNA is bound better than the N-IDR (Figure 7D). Hence, the N-IDR is unlikely to compete effectively with any small RNA, unless the removal of the N-IDR from the RNA-binding cleft requires significant energy. Interestingly, when we disrupted the Pro-Gln interactions described above by mutating Pro(30-33) to Gly(30-33) we found that the N-IDR became more flexible within the RNA-binding groove (Figure S7F-G; Movie S2). Mutation of the whole N-IDR to glycines showed the same effect. However, in both cases N-IDR did not leave the RNA-binding groove as its exit was prevented by a loop (688-695) that covers the RNA-binding groove of WAGO-3 (Figure S7F-G). Substitution of loop residues 688-695 to glycines together with mutation of the whole N-IDR (Residues 1-51) to glycines (Table S1) increased N-IDR flexibility but within simulation time (1 µs) the N-IDR it still stayed within the binding pocket (Figure S7F). Only when we shortened the N-IDR of WAGO-3 with the original amino acid composition Δ(Residues 1-33), did the remaining N-IDR in one of two simulations spontaneously leave the RNA-binding region within 0.1 μs (Figure 7C, S7H), with the distance of residue 34 to helix^2^ increasing to values above 30 Å, while this shortened N-IDR stayed within the RNA binding groove in the other simulation, with the distance to helix^2^ staying below 30 Å. Similarly, in one of two simulations with the N-IDR Δ(Residues 1-29) (Figure 7C), the N-IDR stays in the RNA binding groove in one simulation (distance residue 34-helix^2^ < 30 Å) but moves out of the RNA binding groove in the other simulation (distance residue 34-helix^2^ > 40 Å). These results are consistent with a model in which the removal of an unprocessed N-IDR from the RNA-binding region requires chaperone activities, implying that the described N-IDR conformation could be an effective competitor against 22G RNA binding (Figure 7E).

## DISCUSSION

### Relevance of Argonaute regulation

Argonaute proteins are potent regulators of gene expression. Therefore, cells need to control their activity carefully. Indeed, at least in *C. elegans,* some of the most deleterious phenotypes related to Argonaute proteins are due to erroneous silencing activity, rather than to a simple loss of Argonaute-mediated silencing. For instance, the sterility phenotype that develops over generations in *prg-1* mutants (the worm Piwi homolog) has been attributed to a gain of silencing of histone^57^ and rRNA genes^58,59^. Likewise, when small-RNA inheritance from parents to offspring is compromised, ectopic silencing can induce acute sterility in worms^60,61^. In humans, Argonaute syndrome patients often carry similar missense variants in Argonaute genes^11,12^. Studies of such Argonaute syndrome mutations in *C. elegans* suggest that these variants are not simple loss-of-function but rather neo-morphic alleles^62^. Hence, keeping Argonaute proteins focused on their ‘intended’ targets seems to be crucial, not only in worms but also in humans. The IDR-mediated control over Argonaute loading we uncover here represents a novel mode of Argonaute regulation which prevents mistargeting. We note, that the manuscript by Deng *et al.* (parallel submission) provides independent support for the idea of N-IDR mediated loading control of WAGO proteins. In their study, APP-1 is found enriched in the Mutator foci, which are centres of 22G RNA biogenesis^51^. This spatial arrangement fits well with the role of N-IDR-processing in WAGO-loading.

### IDRs as Argonaute-regulatory entities

In general, IDRs can take regulatory functions by affecting multi-valency, binding affinities, binding specificities and other aspects of the protein function. For instance, auto-regulatory effects of IDRs have been described for kinases and transcription factors^63^. While the regulatory interactions between the IDRs and folded domains are often relatively weak, the fact that they are tethered together within one polypeptide chain increases their effective local concentration, allowing the IDRs to regulate other, higher-affinity interactions of the folded domain. In Argonaute proteins, the N-terminal IDRs function in nuclear localization^18,19^, phase separation^64^, and protein stability regulation^65^ depending on their substantial variations in length and amino acid composition. However, an auto-inhibitory role as we propose here has not yet been described for eukaryotic Argonaute proteins.

A similar regulatory effect of an IDR on a prokaryotic Argonaute system named SPARTA has been described^44^. SPARTA is a heterodimer with a TIR-APAZ subunit and a short prokaryotic Argonaute protein (pAgo) subunit, which is involved in immunity responses in bacteria^66,67^. In the hetero-dimer, the negatively charged C-terminal peptide of the TIR-APAZ subunit is bound in between its own APAZ domain and the MID domain of the unloaded pAgo protein. A regulatory loading function of this arrangement was proposed but not directly demonstrated. Interestingly, the inhibiting C-terminal peptide is negatively charged through the presence of Asp and Glu residues, while the WAGO-1 and WAGO-3 N-IDRs are hardly charged. Also, part of the TIR-APAZ C-terminus adopts an alpha-helical conformation. Even though these specifics are different from what we observed in the WAGO proteins, the underlying concept may be very similar: regulation of small guide RNA binding by IDR-RNA-binding-groove-interactions.

Can this be a more broadly used concept in Argonaute biology? Recent work on Argonaute proteins from *Arabidopsis* and humans has shown that the region directly following the N-IDR responds to loading^68^. This region, also named the N-coil, is accessible to antibodies in an unloaded state, but becomes shielded when the Argonaute is loaded. Even though this does not reveal what the N-IDR is doing during the loading process, it does show that the very N-terminal regions of Argonaute proteins in general are closely involved with the loading process. Further investigation will shed light onto what extent the principle of autoregulation that we uncovered is applicable to other Argonautes. Even if autoregulation were relatively rare, *in-trans* IDR interactions in Argonaute loading processes may not be, as already exemplified by the bacterial SPARTA system. It is worth noting that mammalian Dicer, a key factor in siRNA biogenesis, has long IDR loops, that have not been functionally characterised yet.

Our work supports a model where binding of the N-IDR within the RNA-binding groove of WAGO-1 and WAGO-3 effectively counteracts the loading of small RNAs (Figure 7E). However, alternative modes of regulation are also possible. For instance, in *D. melanogaster,* plants and humans heat shock factors like HSP70 and HSP90 have been shown to modulate the Argonaute loading process^69–72^ and it is possible that the WAGO N-IDRs studied here interact with such factors to mediate the observed effects. Regardless of such alternative possibilities, DPF-3 plays a role in relieving the blocking role of the N-IDRs, and it does so in a non-catalytic manner.

### Catalysis-independent regulation of WAGO-3 by DPF-3

We report catalysis independent effects of the DPF-3 protease on the function of its substrate WAGO-3. This is based on the use of a WAGO-3 N-IDR that is refractory to DPF-3 activity (WAGO-3(P2A;A3E)). The fact that this processing-resilient N-IDR of WAGO-3 does not phenocopy a lack of the N-IDR (*i.e.* sterility) suggests that it can still act as an antagonist to erroneous WAGO-3 loading. WAGO-3 mutant protein with this non-processible N-IDR still needs DPF-3 to localize to PEI granules, and *vice versa* it can still support the recruitment of DPF-3 to these same granules. We also show that this processing mutant of WAGO-3 is normally loaded with small RNAs. All these findings imply that DPF-3 does not need to process WAGO-3 to help loading it. At the same time, loss of catalytic DPF-3 activity by mutation of its active centre does not phenocopy a block in processing by N-IDR mutation. How can this be explained? We propose that these differences come from the trapping of catalytically inactive DPF-3 on its substrates, which extend beyond the WAGO proteins. Effectively, this creates a loss of DPF-3, leading to phenotypes similar to *dpf-3* deletion. Strikingly, mutating the N-IDR of only one of its substrates (WAGO-1) already affects WAGO-3 localization, indicating how potent such trapping of DPF-3 enzyme may be.

### Other catalysis-independent examples of DPF-3 and DPF-3-like enzymes

Activity-independent roles of DPF-3 in *C. elegans* and its human homologs have been reported before. In *C. elegans*, loss of DPF-3 protein, but not DPF-3 activity was shown to affect miRNA function^39^. In particular, animals that expressed a mutant allele of the miRNA-binding Argonaute protein ALG-1 were shown to re-gain miRNA function through loss of DPF-3, caused by increased levels of the ALG-1 paralog ALG-2. How this interplay works, however, remained unclear. Our results provide a potential explanation. First, DPF-3 was shown to physically interact with ALG-1, but not with ALG-2. It is feasible that this interaction helps ALG-1 to be loaded, just like DPF-3 stimulates WAGO-3 loading. The precise nature of the *alg-1* mutant allele used by Harvey et al.^39^ is not clear, but likely it is not a null-allele. Part of the phenotype it drives may therefore stem from mutant ALG-1 protein binding to miRNAs, but then not being (fully) active. If loss of DPF-3 protein would lead to a loss of ALG-1 loading, such unproductive loading into mutant ALG-1 protein would be suppressed, possibly favouring the loading of ALG-2, which subsequently may compensate loss of ALG-1 function.

In human cells, DPP9 (a human DPF-3 homolog) regulates NLRP1 and CARD8 activity in the inflammasome response^43^. It binds to these proteins and keeps them inactive independent of its protease activity. When this interaction is disrupted, NLRP1 and CARD8 can be activated leading to inflammasome formation. There are several intriguing parallels to the mechanism we propose. First, both DPP9 and DPF-3 play a role in controlling the activation of factors related to some form of innate immunity, as the nematode small RNA mechanisms bear many features of the mammalian innate immune responses. Second, this control is, at least partly, catalysis independent. Third, the human homolog of the nematode APP-1, XPNPEP1, has been suggested to act together with DPP9^42^. These similarities may hint at a more deeply conserved role of DPP8-DPF-3 related enzymes in the control of immune-related reactions.

### The role of IDR processing

DPF-3 and APP-1 together process the IDRs of WAGO-1 and WAGO-3. However, the precise role of this post-translational modification of these Argonaute proteins remains unclear. While WAGO processing does not affect loading or subcellular localization, it does affect RNAi inheritance. We can envision that the processing serves as a quality control mechanism at a later step in its activity path. Possibly, unprocessed WAGO-3 protein is unstable upon entry in the oocyte. In this scenario the N-terminal tail may act as a degron, which may be shielded by the presence of DPF-3 in the adult germline, but may become exposed in the zygote. This hypothesis is consistent with N-IDR-sensitive E3 ligases presented by Deng *et al.* (parallel submission). An alternative, non-mutually exclusive possibility is that the N-terminal tail may interfere with inter-molecular interactions that WAGO-3 may need to engage in, in order to trigger an RNAi response. Whatever the precise reason behind the inactivity of unprocessed WAGO-3 in inheritance, this mechanism may play a role in selecting only fully matured WAGO-3 protein to act in inheritance.

Another curious aspect of the N-IDR processing reaction is that WAGO-3, and to a lesser extend WAGO-1, has a number of positions that are intrinsically poor DPF-3 substrates. This already starts at the very first processing step in case of WAGO-3, together with four to five additional poor processing sites along its tail. We anticipate that these sub-optimal processing sites play a role in WAGO-3–DPF-3 interaction, possibly by extending the contact time between DPF-3 and the N-terminal end of WAGO-3. Full WAGO-3 processing can then occur only when local concentration of APP-1 and DPF-3 enzymes are sufficiently high, as would be expect within condensates such as the Mutator foci or PEI granules. It is also conceivable that processing through these sub-optimal sequences is performed or stimulated by additional proteins.

While the manuscript was under development, a recent publication reported a previously unrecognized tripeptidyl peptidase activity for DPF-3^73^. In our studies, we did not detect convincing evidence for DPF-3 tripeptidyl activity *in vitro*. Furthermore, deletion of *app-1* clearly stalled WAGO-1 N-terminal processing after the initial cleavage *in vivo*, whereas processing in the *dpf-3*; *app-1* double deletion was completely blocked. Two potential reasons may explain these observations. First, Xaa-Pro-Pro may be a poor substrate for both the canonical, as suggested here, and tripeptidyl DPF-3 activities. Indeed, His-Pro-Pro exhibited slower tripeptidyl processing efficiency compared to Ser-Gly-Pro under the same reaction conditions^73^. Second, the dipeptidyl activity is generally more efficient than the tripeptidyl activity^73^, making the latter less relevant in the presence of APP-1.

### Limitations of the study

Our work provides several lines of evidence that the N-IDRs of WAGO-1 and WAGO-3 are involved in the regulation of these two Argonautes, in particular to their loading with small RNAs. The IDRs seem to have little effect, if any, on their sub-cellular localisation. Yet, we want to emphasize that the mode in which these IDRs regulate sRNA binding needs further study. First and foremost, we lack direct biochemical evidence that the IDR can be bound in the RNA-binding groove when WAGO-1/3 are not loaded. Another aspect that remains open to interpretation is how DPF-3 affects WAGO function. While we uncovered a processing-independent role in loading, and a processing-dependent role in WAGO-3 function in RNAi inheritance, the above-discussed possibilities and similarities to other systems will need to be tested experimentally. Finally, some of the work in this manuscript depends on tagging proteins with (fluorescent) tags. It cannot be excluded that these tags affect function and/or localization, even if specific functionality assays do not report a phenotype.

## MATERIALS AND METHODS

### Worm culture

*C. elegans* strains were maintained on nematode growth medium (NGM) agar plates seeded with *E. coli* OP50 at 20°C under standard laboratory conditions^74^ unless stated otherwise. Strains expressing WAGO-3 and WAGO-1 with intrinsically disordered region deletion (ΔIDR) were cultured on NGM plates supplemented with 1 mM auxin (indole-3-acetic acid; Alfa Aesar, A10556) to prevent sterility^75^. To induce protein accumulation, gravid adult hermaphrodites with fully developed germlines were collected from auxin-containing plates by washing with M9 buffer. Following two additional M9 washes to remove residual auxin, animals were transferred to NGM plates lacking auxin and incubated for 4 hours prior to RNA immunoprecipitation experiments or for 24 hours before microscopy analysis. The Bristol N2 strain served as the wild-type reference throughout this study. A complete list of strains is provided in the Key Resources Table.

### CRISPR-Cas9 genome editing

For plasmid-encoded Cas9 injections, protospacer sequences were identified using CRISPOR (http://crispor.tefor.net)^76^ and inserted into pRK2411 (a derivative of pDD162, Addgene #47549) through site-directed, ligase-independent mutagenesis (SLIM)^77^. SLIM products were introduced into DH5α competent cells (Invitrogen™) and grown on LB agar containing 100 μg/ml ampicillin. We injected 50 ng/µl Cas9 + co-conversion sgRNA plasmid, 50 ng/µl gene of interest sgRNA plasmid, 750 nM co-conversion ssODN donor (IDT) and 750 nM gene of interest ssODN donor (IDT)^37^.

For Cas9 protein injections, crRNAs were design using the Integrated DNA Technologies CRISPR–Cas9 platform (https://eu.idtdna.com/site/order/designtool/index/CRISPR_CUSTOM). Strains were created by microinjecting 2.5µg/µl recombinant Cas9 protein (produced in-house), 500nM of ssDNA (IDT) or PCR product as repair template, 440 nM co-conversion ssODN donor (IDT), 42.5µM tracrRNA (IDT), 12µM dpy-10 guide RNA (IDT) and 30µM targeted gene guide RNA (IDT) as described previously^78^.

Genome editing used *dpy-10(cn64)* co-conversion approaches^79^. For IDR deletion experiments, injected animals were transferred directly to auxin-supplemented plates. Guide RNA sequences and repair templates are provided in the Key Resources Table. F1 offspring were identified by PCR and validated through Sanger sequencing. Mutants were outcrossed twice with wild-type N2 animals to eliminate potential off-target mutations.

### Male enrichment

Strains carrying *him-5(e1490)* were cultured on egg plates^80^. Male-enriched populations were obtained using a size-based filtration method adapted from Cailloce et al.^81^. Worms were synchronized by alkaline hypochlorite treatment and embryos were hatched overnight in M9 buffer. Synchronized L1 larvae were plated onto 20-24 egg plates (90 mm diameter) and cultured at 20°C for 4 days until reaching one-day adult stage.

Adult worms were collected by rinsing plates with M9 buffer and allowing animals to sediment by gravity at room temperature until a clear pellet appeared, which selectively pelleted adults while retaining most L1 larvae and embryos in suspension. Sedimentation was repeated until the supernatant was visibly clear of small larvae and debris. Approximately 10 mL of settled worms was used to purify males.

The worm suspension was passed through 20 μm cell strainers (pluriSelect, 43-50020-50) (≤2 mL per filter) to retain adults while allowing L1-L3 larvae to pass through. Filters were washed with M9 buffer to remove residual larvae. Adults retained on the filters were pooled and transferred to a 35 μm nylon mesh (Thermofisher Scientific, 12934257) suspended in a beaker filled with M9 buffer. Under these conditions, males passed through the mesh while hermaphrodites were retained. The separation proceeded for approximately 1 hour at room temperature with periodic gentle agitation by pipetting to facilitate male migration through the mesh.

The male-enriched suspension was collected from the beaker and passed through a 30 μm cell strainer (pluriSelect, 43-50030-50) with M9 wash to remove residual embryos and L1 larvae while retaining adult males. Males were recovered from the filter by rinsing with M9 buffer into a 50 mL conical tube. A small aliquot was examined microscopically to assess purity; populations typically contained >90% males. Males were pelleted by centrifugation at 4°C, immediately aliquoted in 200 μl, and flash-frozen in liquid nitrogen for storage at −80°C.

### Determination of N-termini *in vivo*

#### Male native protein extraction

Worms were cultured and filtered to enrich males as described above. Male worm samples were thawed on ice. For each sample, 100 μl of worm suspension was combined with 100 μl of lysis buffer supplied by the TrapR Argonaute Enrichment Kit (Lexogen, 128.24), supplemented with 1× Halt™ Phosphatase Inhibitor Single-Use Cocktail (Thermo Scientific, 78428). After mixing, 150 μl of the lysate was transferred to a fresh microcentrifuge tube and diluted with an additional 250 μl of lysis buffer. The remaining lysate was snap-frozen in liquid nitrogen and stored at −80°C as backup material.

Samples were mechanically disrupted using a BioRuptor (Diagenode, B01020001) sonication device with 10 cycles of 30 s ON/30 s OFF at high intensity at 4°C. Following sonication, lysates were clarified by centrifugation at 20,000 × g for 10 min at 4°C, and supernatants were transferred to new tubes. Total protein concentration was determined using the Pierce BCA Protein Assay Kit (Thermo Fisher Scientific, A65453) according to the manufacturer’s instructions. Protein input was normalized across samples using lysis buffer, with approximately 614 μg of total protein per sample used for subsequent enrichment.

#### Argonaute protein enrichment

Argonaute proteins were enriched using the TrapR Argonaute Enrichment Kit (Lexogen, 128.24) following the manufacturer’s protocol. All steps were performed at room temperature, and kit components were equilibrated to room temperature prior to use. The resin was resuspended by vortexing, and storage buffer was removed by centrifuging at 1,000 × g for 30 s in a 2-ml collection tube. The flow-through was discarded.

250 μl of clarified lysate was applied, mixed by gentle inversion, then centrifuged at 1,000 × g for 30 s. The flow-through was discarded. Bound proteins were eluted by applying 250 μl of TrapR elution buffer and centrifuging into a fresh 2-ml collection tube. A second elution was performed with an additional 250 μl of elution buffer collected into the same tube, yielding a total eluate volume of 500 μl.

Proteins in the 500 μl eluate were precipitated by transferring to a 2-ml microcentrifuge tube and adding 1,100 μl of ultrapure water, followed by 400 μl of TCA stock solution to achieve a final TCA concentration of 20% (v/v). The mixture was vortexed briefly and incubated overnight at 4°C.

Following incubation, samples were centrifuged at 20,000 × g for 30 min at 4°C. The supernatant was carefully removed without disturbing the protein pellet. The pellet was then washed three times with 2 ml of ice-cold 90% (v/v) ethanol, with centrifugation at 20,000 × g for 30 min at 4°C between each wash. After the final wash, the pellet was air-dried at room temperature prior to downstream analysis.

#### Enzymatic protein digestion

The protein pellet was reconstituted in 8 M urea containing 50 mM Na_2_HPO_4_ and 50 mM NaH_2_PO_4_. The urea concentration was reduced to 6 M using ammonium bicarbonate buffer and to a final concentration of 50 mM ammonium bicarbonate. Proteins were further processed using the SP3 method ^82^. Proteins were reduced in 5 mM DTT at room temperature, alkylated in 15 mM iodoacetamide in the dark and quenched in 5 mM DTT. Enzymatic protein digestion was performed using trypsin at 37°C overnight. Following acidification by formic acid, the peptides were purified by solid phase extraction in a C18 StageTip format^83^.

#### Liquid chromatography tandem mass spectrometry

Peptides were separated via an in-house packed 45-cm analytical column (inner diameter: 75 μm; ReproSil-Pur 120 C18-AQ 1.9-μm silica particles, Dr. Maisch GmbH) on a Vanquish Neo UHPLC system (Thermo Fisher Scientific). Online reversed-phase chromatography was performed through a 70-min non-linear gradient of 1.6-48% acetonitrile with 0.1% formic acid at a nanoflow rate of 300 nl/min. The eluted peptides were sprayed directly by electrospray ionization into an Orbitrap Astral mass spectrometer (Thermo Fisher Scientific). Mass spectrometry measurement was conducted in data-dependent acquisition mode using a top50 method with one full scan in the Orbitrap analyzer (scan range: 325 to 1,400 m/z; resolution: 120,000, target value: 3 × 106, maximum injection time: 25 ms) followed by 50 fragment scans in the Astral analyzer via higher energy collision dissociation (HCD; normalized collision energy: 26%, scan range: 150 to 2,000 m/z, target value: 1 × 104, maximum injection time: 10 ms, isolation window: 1.4 m/z). Precursor ions of unassigned, +1 or higher than +7 charge state were rejected. Additionally, precursor ions already isolated for fragmentation were dynamically excluded for 15 s.

#### Mass spectrometry data analysis

Raw data files were processed by MaxQuant software (version 2.6.2.0)^84^ using its built-in Andromeda search engine^85^. MS/MS spectra were searched against a target-decoy database containing the forward and reverse protein sequences of the WormPep release WS292 (28,587 entries), the UniProt E. coli K-12 reference proteome (release 2022_03; 4,460 entries), a list of N-terminally truncated WAGO-1/WAGO-3 proteins, WAGO-3 (P2G, A3E) mutant and a default list of common contaminants. Trypsin/P specificity was assigned. Carbamidomethylation of cysteine was set as fixed modification. Methionine oxidation and protein N-terminal acetylation were chosen as variable modifications. A maximum of 2 missed cleavages were allowed. The “second peptides” option was switched on. “Match between runs” was activated. The minimum peptide length was set to 7 amino acids. The maximum peptide mass was adjusted to 6,500 Da. False discovery rate (FDR) was set to 1% at both peptide and protein levels.

Peptide intensities were normalized by median-centering on the logarithmic scale. N-terminally truncated tryptic peptides of WAGO-1/WAGO-3 detected in at least 3 out of the 4 replicates from at least one of the five strains were retained. These peptide intensities were then displayed on a heat map.

### Determination of N-termini in vitro

#### Recombinant Protein Purification

Purification of *C. elegans* APP-1 WT and the catalytic-dead mutant (D413Q, D424N) was performed as described^46^. Briefly, 6xHis-tagged APP-1 WT and (D413Q, D424N) mutant were cloned into the pET-His6 (pET-28a) vector and transformed into *Escherichia coli* Arctic BL21 (DE3) cells. Cultures (500 mL) were grown at 37°C until OD600 reached ∼0.6, after which expression was induced by adding 1 mM isopropyl β-D-1-thiogalactopyranoside (IPTG) and incubated overnight at 16°C. Cells were harvested by centrifugation at 4,000 × g for 20 min, and pellets were resuspended in 10 pellet volumes of lysis buffer (10 mM Tris-HCl pH 8.0, 300 mM NaCl, 10% glycerol (v/v)), supplemented with 0.2% Triton X-100, 20 mM imidazole, and cOmplete EDTA-free protease inhibitor cocktail tablets (Roche, REF 11836170001). Cells were lysed on ice using a Branson Sonifier 450 (9 mm tip, output 6, 30% duty cycle, 3 min). Lysates were cleared by centrifugation at 40,000 × g for 20 min at 4°C. Cleared lysates were loaded onto a 5-mL HiTrap column (Cytiva) and washed with 20 column volumes of lysis buffer supplemented with 20 mM imidazole, followed by elution using a linear gradient from 10–300 mM imidazole in wash buffer over 10 column volumes. The eluted peak was concentrated, injected using a 5-mL loop, and further purified over a Superdex 200 size-exclusion chromatography column (Cytiva), pre-equilibrated with storage buffer (10 mM Tris-HCl pH 8.0, 150 mM NaCl, and 10% glycerol (v/v)). All purification steps were performed on an automated chromatography system (Cytiva, ÄKTA go™). The purified APP-1 proteins were concentrated to 5–10 mg/mL, aliquoted, snap-frozen, and stored at −80°C.

Purification of *C. elegans* DPF-3 WT and catalytic-dead mutant (D861A) was performed as described^38^, with modifications. Both proteins carried an N-terminal 6xHis-MBP-3C–tag and a C-terminal TwinStrep tag were expressed using the Baculovirus-insect cell system (Bac-to-Bac, Thermo Fisher Scientific) and the modified EmBacY genome^86^. *Spodoptera frugiperda* SF9 cultures (2–3 L) were grown in SF900 III serum-free medium (Thermo Fisher Scientific, Cat# 12658019) and infected with passage 1 virus at a density of 1 × 10^^6^ cells/mL. Cells were harvested 50 hours post-infection by centrifugation at 800 × g for 15 min at 4°C, resuspended in 100 mL lysis buffer per liter of culture (50 mM Tris-HCl, 500 mM NaCl, 10 mM imidazole, 2 mM MgCl_2_, 1 mM TCEP, 5% glycerol, pH 8.0), supplemented with 0.1% Triton X-100, 20 mM imidazole, cOmplete EDTA-free protease inhibitor cocktail (Roche, REF 11836170001), and 125 U/mL Benzonase. Cell lysis was performed on ice using a Branson Sonifier 450 (9 mm tip, output 6, 15% duty cycle, 2 min). Lysates were cleared by centrifugation at 40,000 × g for 30 min at 4°C and applied to a 5-mL HisTrap FF column (Cytiva) on an automated chromatography system (Bio-Rad NGC Quest Plus). The column was washed with 20 column volumes of wash buffer (50 mM Tris-HCl, 500 mM NaCl, 10 mM imidazole, 5% glycerol, pH 8.0) and eluted with a linear gradient of 10–500 mM imidazole over 10 column volumes. Eluted peaks were diluted 1:2 in Strep-Tactin wash buffer (50 mM Tris-HCl, 500 mM NaCl, 2 mM DTT, 5% glycerol, pH 8.0), added to 6 mL equilibrated Strep-Tactin 4Flow slurry (IBA-Lifescience), and incubated overnight with 0.5 mg 6xHis-3C protease (in-house produced) at 4°C with rotation to remove the N-terminal 6xHis-MBP tag. Beads were washed twice with Strep-Tactin wash buffer by centrifugation at 500 × g for 5 min at 4°C, then transferred into an Econo-Pac column (Bio-Rad) for three washes in storage buffer (30 mM Hepes-NaOH pH 7.4, 300 mM NaCl, 10% glycerol). DPF-3-twinstrep enzymes were eluted in storage buffer containing 2.5 mM desthiobiotin, concentrated to 10–14 μM, snap-frozen in liquid nitrogen, and stored at −80°C.

#### In Vitro Cleavage Assay and Mass Spectrometry Data Analysis

The assay was modified from Gudipati et al., 2021^38^. Substrate peptides (Key Resources Table) were obtained from GenScript and resuspended in TBS buffer (50 mM Tris-HCl pH 7.5, 150 mM NaCl) with 2 mM TCEP to a final concentration of 0.1 nmol/μL (stock). For each reaction, stock peptide was diluted to 1 μM and combined with 0.1 μM recombinant protein(s) (DPF-3 WT, DPF-3 [D861A), APP-1 WT, or APP-1 [D413Q; D424N]). Reaction volume depended on the number of time points collected, with 50 μL per time point. Assays were performed at 21°C in a ThermoMixer C (Eppendorf); aliquots were withdrawn at indicated times and mixed with 100 μL of 0.5% formic acid to stop the reactions. Samples were snap-frozen and stored at −20°C until mass spectrometry.

Acidified peptides were purified via strong cation exchange (SCX) using StageTip format^83^. Peptides bound to SCX were washed sequentially in 0.1% formic acid and then in 80% acetonitrile with 0.1% formic acid. Peptides were eluted in 80% acetonitrile with 1% ammonia, dried by centrifugal evaporation, and reconstituted in 80% acetonitrile with 0.1% formic acid. Acetonitrile was removed by a second centrifugation before liquid chromatography. Peptides were separated on an EASY-nLC 1200 system (Thermo Scientific) using a 45-cm analytical column (75 μm ID, ReproSil-Pur 120 C18-AQ 1.9-μm beads), applying a 60-min non-linear gradient (1.6–40% acetonitrile, 0.1% formic acid) at 300 nL/min. Eluted peptides were analysed by electrospray ionization on a Q Exactive Plus Orbitrap mass spectrometer (Thermo Scientific), in data-dependent acquisition mode (top10 method) with specified scan and fragmentation parameters. Unassigned or +1 charge precursor ions were rejected and dynamically excluded for 10 s.

Raw MS data were processed with MaxQuant (v2.6.2.0)^84^ using the Andromeda search engine^85^, in “no digestion” mode against a target-decoy database of peptide sequences listed in the Key Resources Table and common contaminants. Methionine oxidation was set as a variable modification (max 4 per peptide). “Second peptides” was enabled, and “match between runs” was disabled. Minimum peptide length was 7 amino acids; maximum mass 5,000 Da. Peptide and protein FDR were set to 1%. Peptide intensities were exported to Excel (Microsoft 365, v16.98), where relative intensities were calculated for each sample to measure the proportion in the population of each peptide. These values were compared across samples and time points. Some WAGO-3 WT replicates were removed due to accumulative intensity being less than 10% with respect to other samples. Comparisons were plotted as barplots using Rstudio (v2025.05.1+513).

### Tc1 Reactivation assay

20 worms carrying the *unc-22(st136::Tc1) IV* allele in either wild-type or mutant backgrounds, displaying the Unc phenotype, were picked onto 20 10-cm OP50 plates per genotype and grown until starvation. Upon starvation, the plates were screened for reversions to the wild-type phenotype and the reversion frequency was estimated using the formula: f = -ln [(T - R) / T] / N, where T = total number of plates scored, R = number of plates with revertants, and N = number of worms on the plate^48^. On starved plates, the number of worms was estimated to be 10.000.

### RNA interference

RNAi feeding experiments using *E. coli* HT115(DE3) expressing double-stranded RNA^87^. Empty vector L4440 (Addgene plasmid #1654; gift from Andrew Fire) served as a negative control. Two RNAi constructs were used: one plasmid targeting the transgene *mjIs31*[*pie-1p::GFP::his-58::pie-1 3’UTR*] II^36^(gift from Scott Kennedy) and pRFK4103 targeting *xfSi255*[*his-67p::gfp::his-67::tbb-2 3’UTR + Cbr-unc-119(+)*]^35^. HT115(DE3) bacteria transformed with each construct were streaked onto LB agar plates supplemented with 100 μg/mL ampicillin and 10 μg/mL tetracycline and incubated overnight at 37°C. Single colonies were inoculated into 50 mL LB broth without antibiotics and cultured overnight at 37°C with shaking. Cultures were then used to seed NGM plates.

### Biparental RNAi inheritance

Strains expressing GFP::H2B fusion proteins in wild-type^36^ (gift from Scott Kennedy) or specified mutant backgrounds were synchronized by alkaline hypochlorite treatment, and embryos were transferred to NGM plates seeded with HT115(DE3) bacteria expressing dsRNA targeting *gfp* or L4440 empty vector control. After 4 days at 20°C, P0 adults were screened for GFP silencing by fluorescence microscopy.

P0 hermaphrodites were treated with alkaline hypochlorite to isolate F1 embryos, which were transferred to NGM plates seeded with OP50 *E. coli* (no RNAi). After 4 days at 20°C, F1 adults were scored for GFP expression. Animals were classified into two categories: “GFP-expressing” (fluorescence intensity comparable to non-RNAi controls) or “silenced” (complete absence of detectable GFP or barely detectable fluorescence).

Fluorescence imaging was performed using a Leica AF7000 widefield microscope equipped with a Hamamatsu sCMOS Flash 4.0 camera and a 40×/1.1 NA water immersion objective. Images were acquired using LASX Application Suite X software (version 3.7.4.23463). Image processing and visualization were performed using ImageJ (version 1.53c).

### Paternal RNAi inheritance

Paternal RNAi inheritance was assessed as described by Schreier et al.^35^. Gravid adult hermaphrodites were treated with alkaline hypochlorite solution to isolate embryos, which were transferred to *gfp* RNAi or control feeding plates. Animals were maintained at 20°C and silencing penetrance was assessed by scoring roughly 50 males per condition at the adult stage.

P0 males exhibiting RNAi-induced silencing were crossed to L4 hermaphrodites that had been maintained on OP50 *E. coli* control plates. Five mating plates were established, each containing 10 P0 males and 5 hermaphrodites on OP50 seeded plates. After 24 hours, individual hermaphrodites from each mating plate were transferred to separate OP50 plates (5 hermaphrodites per condition, each on individual plates) and allowed to lay eggs for 48 hours.

Cross-progeny (F1) were identified by the presence of the mCherry marker from the paternal parent. For quantitative analysis of silencing inheritance, tile scan images of approximately 50 F1 males per condition were acquired at the adult stage using a Leica THUNDER Imager inverted widefield microscope with a 10×/0.32 NA dry objective.

### Western Blot

For Western blot analysis, 100–200 adult worms were manually collected and lysed in 1× NuPAGE™ LDS sample buffer (Invitrogen, NP0007) supplemented with 100 mM DTT, followed by incubation at 95°C for 30 min. For male samples, the males were separated from hermaphrodites one day before collection. Protein lysates were resolved on NuPAGE 3–8% Tris-Acetate gels (Thermo Fisher Scientific, EA0375BOX) using Tris-Acetate SDS running buffer (Thermo Fisher Scientific, LA0041), or NuPAGE 4–12% Bis-Tris gels (Thermo Fisher Scientific, MP0321BOX) using MOPS NuPAGE running buffer (Thermo Fisher Scientific, NP0001), at 120 V, with Prestained Protein Standard (Thermo Fisher Scientific, 26616) included for molecular weight reference.

Separated proteins were electrotransferred onto Amersham™ Protran® Western blotting nitrocellulose membranes (SigmaAldrich, GE10600002), at 140 V for 1 hour using a Transfer buffer (prepared in-house) containing 15% methanol. Membranes were blocked for 1 hour in PBS supplemented with 5% skim milk and 0.1% Tween-20, then cut into sections based on molecular weight markers. Individual membrane sections were incubated overnight at 4°C with primary antibodies diluted in blocking buffer: mouse anti-FLAG (1:2000; Sigma-Aldrich, F3165) and mouse anti-α-tubulin (1:4000; Abcam, ab7291). Following five washes of 5 min each with PBS containing 0.1% Tween-20 (PBS-T), membranes were incubated for 1 hour with IRDye 800CW-conjugated goat anti-mouse secondary antibody (1:10,000; LI-COR, 926-32211), then washed three additional times with PBS-T. Immunoreactive bands were visualized using a LI-COR Odyssey M imaging system. Band intensities were quantified using Empiria Studio v 3.3 and the intensities for proteins of interest were normalized against α-tubulin.

The intensity fold change relative to wild-type was quantitatively analysed on a log-scale. To evaluate potential residual batch effects after normalization, we fitted a hierarchical statistical model (Gaussian mixed effect model) which incorporated the experimental day as a covariate and compared it with a reduced model without batch effect. Comparison of Akaike information criterion (AIC) values and a Likelihood Ratio Test indicated no relevant batch effect. For the analysis an empirical Bayes-moderated paired t-tests as implemented in the limma 3.64.3 R-package^88^ was used.

For the comparisons of fold changes between mutants, contrasts in Gaussian linear models were tested, and the multivariate t-method, as implemented in the emmeans 2.0.0 R-package, was used for multiple testing correction.

### Counting of sterile animals

To assess germline fertility, 30 L1-stage larvae per replicate were transferred to auxin-free NGM plates for each strain. After 4 days of development at 20°C, individual hermaphrodites were scored for the presence of embryos by visual inspection under a dissecting microscope. Animals lacking visible embryos were classified as sterile.

### Immunostaining

Adult animals were dissected in Egg buffer (25 mM HEPES, pH 7.4, 118 mM NaCl, 48 mM KCl, 2 mM EDTA, 0.5 mM EGTA, 10mM NaN_3_) supplemented with 0.1% Tween-20 and fixed in 0.3% formaldehyde diluted in Egg buffer. Following freeze-cracking on dry ice, specimens were transferred to pre-chilled acetone and incubated for 10 min. Germ lines were incubated with anti-GFP nanobody abberior STAR GREEN 1:500 dilution (NanoTag Biotechnologies, N0304-AbGREEN-L), primary anti-HA antibody at 1:1000 dilution (Sigma–Aldrich, H3663), and secondary anti-mouse IgG Alexa Fluor 555 at 1:1000 dilution (Thermo Fisher A11029). Samples were mounted using ibidi mounting medium containing DAPI (ibidi, 50011). Airyscan imaging of immunostained samples was performed on a Zeiss LSM900 microscope with a 63× oil-immersion objective. Images were analysed using ImageJ 1.54p^89^ and Omero^90^.

### Single-molecule FISH

Single-molecule FISH (smFISH) probes targeting the *his-60* RNA (Key Resources Table) with Quasar670 dye were designed using Stellaris Probe Designer (v4.2). For smFISH, worms were collected at 30 h (L2) and 48 h (L3) after L1 plating. Animals were washed twice in M9 buffer and fixed in 4% paraformaldehyde (Sigma-Aldrich, 441244-1KG) in 1× PBS at room temperature for 1 h. Following fixation, worms were centrifuged at 10,000 × *g* and washed with 1 ml 1× PBS. For permeabilization, worms were kept at 4°C overnight in 70% ethanol. The following day, samples were washed once with 1 ml wash buffer (10% (v/v) deionized formamide (Thermo Fisher Scientific, AM9342), 2× SSC) and resuspended in 100 μl hybridization buffer (100 mg/ml dextran sulfate (Sigma-Aldrich, D4911-50G), 1 mg/ml *E. coli* tRNA (Roche, 10109550001), 2 mM vanadyl ribonucleoside complex (NEB BioLabs, S1402S), 0.2 mg/ml BSA (Sigma-Aldrich, A7906-500G), 10% (v/v) deionized formamide (Thermo Fisher Scientific, AM9342)) along with 2.5 μl of 5 μM probe in TE buffer. Samples were incubated overnight at 30°C and washed twice with 1 ml wash buffer at 30°C for 30 min; the second wash included 5 ng/ml DAPI. Samples were then washed twice in 2× SSC buffer and twice in 1× PBS. Finally, animals were resuspended in ibidi mounting medium with DAPI (ibidi, 50011) and placed on µ-slides for microscopy (ibidi, 80606). smFISH samples were imaged on a BC43 spinning disk confocal microscope with a 40×/0.75 air objective. Images were analysed using ImageJ 1.54p^89^ and Omero^90^.

### Live imaging

For live imaging, 20–30 animals were washed in 100 µl M9 buffer and then moved to 50 µl M9 buffer containing 40 mM NaN_3_ (Sigma-Aldrich, S002) on a coverslip. After worms stopped moving, a glass slide with a freshly prepared 2% (w/v) agarose pad in water was placed on top of the coverslip.

For imaging of sperm cells, males were separated from hermaphrodites overnight to avoid recent mating and then dissected in 10 µl sperm buffer (50 mM HEPES pH 7.5, 50 mM NaCl, 25 mM KCl, 5 mM CaCl2, and 1 mM MgSO4, 10 mM glucose) in a µ-Slide 15 Well 3D with glass bottom (Ibidi, 81507). The corpses were removed and another 30 µl sperm buffer carefully added to the well. Confocal imaging was carried out on a Leica Stellaris 8 falcon confocal microscope using a 20x/0.75 dry, 63×/1.4 or 100×1.4 oil-immersion objective. Images were analysed using ImageJ 1.54p^89^ and Omero^90^.

### Image processing and quantification

To quantify paternal RNAi inheritance of GFP silencing, tile scan images were analysed using a custom ImageJ macro (Fiji version 1.53c). To eliminate edge artifacts from tile stitching, a 100-pixel border was cropped from all sides of each image prior to analysis.

Individual worms were segmented using fluorescence signals from the RFP channel. The segmentation workflow consisted of the following steps: (1) Gaussian blur (σ = 4 pixels) was applied to reduce noise, followed by intensity thresholding (0-150 arbitrary units) to generate a binary mask; (2) the mask was further refined with an additional Gaussian blur (σ = 2 pixels) and thresholding step to improve worm boundary detection; (3) particle analysis was performed to detect objects within size (2,500-9,000 pixels²) and circularity (0.00-0.75) constraints, which effectively excluded debris and overlapping worm clusters.

For each image, a background region of interest (ROI) was defined as the inverse of all detected bright objects. Mean background intensity was calculated from this ROI and subtracted from the entire image for both the GFP channel and the RFP channel independently. This correction normalized for variations in illumination and background autofluorescence across tile scans.

Following automated segmentation, ROIs were manually inspected and any erroneously detected non-worm objects were removed. For each validated worm ROI, integrated fluorescence intensity and mean intensity were measured in both channels. Area, mean intensity, and integrated density values were exported to separate results tables for each channel for subsequent statistical analysis.

It was expected that the integrated GFP intensity would be biased by individual size, and the mean intensity by positional effects. To correct both biases simultaneously, the ratio of integrated GFP and RFP intensities was calculated :

RFI = RawIntDen(GFP) / RawIntDen(RFP).

Note that calculating a ratio image with the microscopy software is not advisable due to systematic differences in the morphological structure of the GFP and RFP signals.

The RFI response at log_2_ scale was analysed using Gaussian linear models with categorical predictors *genotype* and *paternal RNAi treatment* (including their interaction). The normality of log_2_(RFI) values was assessed via Q-Q plots and Shapiro-Wilk tests. For the P0 data set in Fig. 2D, the distributional assumptions were borderline. However, given the large sample sizes (n > 50 per group), analysis with Gaussian models remained appropriate. Because group variances differed, inverse-variance weighting was applied. For the sample size at hand, this is an efficient method for correcting bias in standard-error estimators caused by heteroskedasticity.

Error bars in Fig. 2C-E represent 95% confidence intervals for the group means, with the end points transformed to original scale. The *p*-values were corrected for multiple testing within each contrast type (RNAi-treated vs. control across genotypes; mutants vs. wild-type in RNAi treated group) using the multivariate t-method implemented in emmeans.

Models were fitted using the *lm* function in R 4.5.2, and contrast *p*-values and confidence intervals were calculated with the emmeans 2.0.0 package.

For smFISH analyses, confocal z-stack projections (sum of slices projections of 13 optical sections spanning 6 μm per animal) were used. For each animal, a single gonad arm was selected for analysis, with regions of interest (ROIs) delineated based on PGL-1::mTagRFP-T fluorescence, while DAPI staining was used to define whole-animal ROIs. Background ROIs were established by inverting the whole-animal masks. These ROIs were applied to the red (mTagRFP-T) and far-red (*his-60* smFISH) channels of Sum of slices projections to extract the following parameters: area, mean intensity, standard deviation, mode, minimum, and maximum values. Each replicate corresponds to an independently treated animal, constituting biological replicates.

To calculate normalized *his-60* smFISH signals, the background-corrected mean intensities for both *his-60* smFISH (gonad mean intensity − background mean intensity) and PGL-1::mTagRFP-T (gonad mean intensity − background mean intensity) were first determined. The normalized *his-60* signal was then computed as the ratio of background-corrected *his-60* intensity to background-corrected PGL-1::mTagRFP-T intensity. Statistical significance was assessed using Welch’s t-test for WAGO-1 experiments and Dunnett’s multiple comparison test for WAGO-3 experiments.

### RNA immunoprecipitation

Synchronized animals were maintained at 20°C on auxin-supplemented plates until reaching the young adult stage. Worms were washed three times with M9 buffer, transferred to auxin-free plates, and incubated for 4 hours before collection. Animals were then washed with M9 buffer, resuspended in sterile water as 300 μl aliquots, and flash-frozen on dry ice.

For RNA immunoprecipitation (RIP), frozen aliquots were thawed on ice and combined 1:1 (v/v) with 2× lysis buffer containing 50 mM Tris-HCl (pH 7.5), 300 mM NaCl, 3 mM MgCl₂, 2 mM dithiothreitol (DTT), 0.2% Triton X-100, cOmplete Mini EDTA-free Protease Inhibitor Cocktail (Roche, 11836170001), and 160 U/μl RNase Inhibitor, Murine (New England BioLabs, M0314). Lysates were sonicated using a Bioruptor Plus (Diagenode, B01020001) with 10 cycles of 30 s on/30 s off at 4°C, followed by centrifugation at 21,000×*g* for 10 min at 4°C. A 100 μl supernatant aliquot from each sample was reserved as input material for subsequent RNA extraction.

For RIP, 30 μl Anti-FLAG M2 Magnetic Beads (Sigma-Aldrich, M8823) were equilibrated with 500 μl 1× wash buffer (25 mM Tris-HCl pH 7.5, 150 mM NaCl, 1.5 mM MgCl₂, 1 mM DTT, cOmplete Mini EDTA-free Protease Inhibitor Cocktail, 80 U/μl RNase Inhibitor, Murine), then incubated with the remaining lysate for 2 hours at 4°C with rotation. Following three washes with 500 μl 1× wash buffer, RNA from both input and RIP samples was extracted using TRIzol Reagent (Invitrogen, 15596018) combined with the Direct-zol RNA kit (Zymo Research, R2050) according to manufacturer protocols. RNA quality and quantity were assessed using the Bioanalyzer RNA 6000 Nano Kit (Agilent Technologies, 5067-1511) and Qubit RNA BR Assay Kit (Invitrogen, Q10210), respectively.

### Sample preparation for sequencing of total small RNA populations

For sequencing of total small RNA populations, worms were expanded on 10 NGM plates seeded with OP50 *E. coli*. Gravid adults were treated with alkaline hypochlorite solution, and isolated embryos were resuspended in M9 buffer and incubated at 20°C for 24 hours to synchronize hatching. Synchronized L1 larvae were plated onto 25 NGM plates per strain and cultured at 20°C. Worms were harvested at the gravid adult stage (approximately 63 hours post-plating).

Worms were collected by rinsing plates with ice-cold M9 buffer and pelleted by centrifugation at 4°C. The washing step was repeated three times with M9 buffer, followed by three washes with ultrapure water to remove residual bacteria and salts. Excess liquid was removed, and 50 μL packed worm pellets were transferred to low-bind microcentrifuge tubes. Samples were flash-frozen in liquid nitrogen and stored at −80°C until RNA extraction.

Worm pellets (50 μL) were resuspended in 500 μL TRIzol LS Reagent (Invitrogen, 10296028) and mixed thoroughly by pipetting and vortexing. To disrupt worm cuticles, samples underwent six freeze-thaw cycles consisting of 30 seconds in liquid nitrogen followed by thawing in a 37°C water bath for approximately 2 minutes with vortexing between cycles. Complete lysis was confirmed by microscopic examination; additional freeze-thaw cycles were performed if intact worms remained visible.

Lysates were centrifuged at 21,000 × g for 5 minutes at room temperature to pellet debris. The supernatant (approximately 500 μL) was carefully transferred to a fresh tube, avoiding the upper lipid layer and debris pellet. The supernatant was mixed 1:1 (v/v) with 100% ethanol (550 μL) and processed immediately using the Direct-zol RNA Miniprep Kit (Zymo Research, R2050) according to the manufacturer’s protocol.

### Small RNA sequencing

RppH (NEB) treatment was performed with a starting amount of 100-200ng. After purification samples were quantified using the Qubit RNA High Sensitivity Assay Kit. NGS library prep was performed with NEXTflex Small RNA-Seq Kit V3 with UDIs following Step A to Step G of PerkinElmer’s standard protocol (V19.09) using the NEXTFlex 3’ SR Adaptor and 5’ SR Adaptor (5’rApp/NNNNTGGAATTCTCGGGTGCCAAGG/3ddC/and5’GUUCAGAGUUCUACAGUCC GACGAUCNNNN, respectively). Step A (NEXTflex 3’ 4N Adenylated Adapter Ligation) was performed overnight at 20°C. Libraries were prepared with a starting amount of 45 ng and amplified in 23 PCR cycles.

Amplified libraries were pooled in equimolar ratio and the pool was purified by running a 3% agarose gel cassette on a PippinHT (sage science) and size-selected for bands in 152-200 bp size range (adapters + insert).

The Pool was profiled in a High Sensitivity DNA Chip on a 2100 Bioanalyzer (Agilent technologies) and quantified using the Qubit 1x dsDNA HS Assay Kit in a Qubit 4.0 Fluorometer (Life technologies).

The pool was sequenced on 1 NextSeq 2000 P1 Flowcell, SR for 1x 84 cycles plus 2x 8 cycles for the Dual Index Read. For the sequencing of the i5 Barcode a custom sequencing i5 primer (NextFlex) was used.

Sequencing data from RIP-Seq libraries^24^ were analysed using the tinyRNA pipeline^91^, with read alignment performed against the C. elegans WormBase WS279 reference genome. Small RNA sequencing data from mutant and wild type libraries were processed using Cutadapt (https://doi.org/10.14806/ej.17.1.200) for adapter removal (-a TGGAATTCTCGGGTGCCAAGG -O 5 -m 26 -M 48) and low-quality reads were filtered out using the FASTX-Toolkit (fastq_quality_filter, -q 20 -p 100 -Q 33). Unique molecule identifiers were used to remove PCR duplicates using a custom script and were subsequently removed using seqtk (trimfq-l 4 – b 4). Reads were aligned using bowtie (v.1.3.1)^92^ (--phred33-quals -- tryhard --best --strata –chunkmbs 256 -v 2 -M 1) and 22G-RNAs (defined as any read of length 20-23 nt with no bias at the 5’-position) were extracted using a python script available at https://github.com/adomingues/filterReads. Reads were counted using htseq-count (v.2.0.2)^93^ (-s no -m intersection-nonempty), and targets were determined using DeSeq2^94^ in R, with a p-value cutoff of 0.01 and a cutoff for log2(fold change) of 1. Sequencing data from a *dpf-3* mutant was downloaded from GEO (accession: GSE151717) and processed as described^38^. Targets were determined by DeSeq2, using the same parameters as for the other deletion mutants. Data visualization was conducted in R.

### Immunoprecipitation followed by mass spectrometry

#### Immunoprecipitation of DPF-3

Worms were cultured and filtered to enrich males as described above (Male enrichment section). Frozen worm samples were thawed on ice and resuspended in 500 μL of cold lysis buffer (25 mM Tris-HCl pH 7.5, 150 mM NaCl, 1.5 mM MgCl₂, 1 mM DTT, 0.1% Triton X-100, and cOmplete Mini EDTA-free protease inhibitor cocktail (Roche, 11836170001). Lysates were sonicated for 10 cycles of 30 seconds on and 30 seconds off at high efficiency, with sonication repeated if cells were not completely lysed. Samples were centrifuged at maximum speed for 10 minutes at 4°C, and supernatants were collected. Protein concentrations were determined by BCA assay, and equal amounts of protein were used for each sample. Input samples (13 μL) were removed and stored at −20°C.

For immunoprecipitation, 20 μL of Chromotek GFP-TRAP magnetic agarose beads (Proteintech, 17323363) were washed three times with 500 μL of wash buffer for 10 minutes each at 4°C with rotation. After the final wash, 450 μL of lysate was added to the beads. The mixture was incubated overnight at 4°C with rotation and 13 μL of flow-through was collected and stored at −20°C. Beads were then washed three times with 500 μL of wash buffer for 10 minutes each at 4°C with rotation. Proteins were eluted by adding 50 μL of resuspension buffer (25 mM Tris-HCl pH 8.0, 1% SDS) and boiling at 70°C for 10 minutes. Eluates were collected and stored at −20°C until analysis.

#### Enzymatic protein digestion

The eluted proteins were processed using the SP3 approach^82^. Proteins were reduced in 5 mM DTT, alkylated in 15 mM iodoacetamide in the dark and quenched in 5 mM DTT. Enzymatic protein digestion was performed using trypsin at 37°C overnight. Following acidification by formic acid, the peptides were purified by solid phase extraction in a C_18_ StageTip format^83^.

#### Liquid chromatography tandem mass spectrometry

Peptides were separated via an in-house packed 45-cm analytical column (inner diameter: 75 μm; ReproSil-Pur 120 C_18_-AQ 1.9-μm silica particles, Dr. Maisch GmbH) on a Vanquish Neo UHPLC system (Thermo Fisher Scientific). Online reversed-phase chromatography was performed through a 70-min non-linear gradient of 1.6-32% acetonitrile with 0.1% formic acid at a nanoflow rate of 300 nl/min. The eluted peptides were sprayed directly by electrospray ionization into an Orbitrap Astral mass spectrometer (Thermo Fisher Scientific). Mass spectrometry measurement was conducted in data-dependent acquisition mode using a top50 method with one full scan in the Orbitrap analyser (scan range: 325 to 1,300 m/z; resolution: 120,000, target value: 3 × 10^6^, maximum injection time: 25 ms) followed by 50 fragment scans in the Astral analyser via higher energy collision dissociation (HCD; normalized collision energy: 26%, scan range: 150 to 2,000 m/z, target value: 1 × 10^4^, maximum injection time: 10 ms, isolation window: 1.4 m/z). Precursor ions of unassigned, +1 or higher than +6 charge state were rejected. Additionally, precursor ions already isolated for fragmentation were dynamically excluded for 15 s.

#### Mass spectrometry data analysis

Raw data files were processed by MaxQuant software (version 2.6.2.0)^84^ using its built-in Andromeda search engine^85^. MS/MS spectra were searched against a target-decoy database containing the forward and reverse protein sequences of the WormPep release WS292 (28,587 entries), the UniProt *E. coli* K-12 reference proteome (release 2022_03; 4,460 entries), the transgenic GFP tag and a default list of common contaminants. Trypsin/P specificity was assigned. Carbamidomethylation of cysteine was set as fixed modification. Methionine oxidation and protein N-terminal acetylation were chosen as variable modifications. A maximum of 2 missed cleavages were allowed. The “second peptides” option was switched on. “Match between runs” was activated. The minimum peptide length was set to 7 amino acids. The maximum peptide mass was adjusted to 6,500 Da. False discovery rate (FDR) was set to 1% at both peptide and protein levels.

The MaxLFQ algorithm^95^ was employed for label-free protein quantification without using its default normalization option. Minimum LFQ ratio count was set to one. Both the unique and razor peptides were used for quantification. The LFQ intensities were normalized by median-centering on the logarithmic scale. Reverse hits, potential contaminants and “only identified by site” protein groups were first filtered out. Proteins were further filtered to retain only those detected in at least three out of the four replicates in either one of the two strains in each comparison. Following imputation of the missing LFQ intensity values, a linear model was fitted using the limma package in R^96^ to assess the difference between the two strains for each protein. The log_2_ fold change and the significance of the difference were displayed on a volcano plot. Only proteins with a minimum log_2_ fold change of 1 and a *p* value lower than 0.01 were considered differentially regulated.

### RNA loading assay

The immunoprecipitated WAGO-3 RNA was obtained following the RNA immunoprecipitation from male-enriched worm suspension procedure, as described in the RNA immunoprecipitation and Determination of N-termini in vivo sections. An aliquot of the native protein extract (10%) was frozen as input control for western blot. The 5′ triphosphate from the immunoprecipitated RNA was removed by treatment with Quick CIP (NEB, Cat. #M0525) to eliminate any phosphates present at the 5′ ends of small RNAs. In a total reaction volume of 15 µL, 10 µL of RNA was incubated with 1× rCutSmart™ Buffer (NEB, Cat. #B6004S), 5 units of Quick CIP (NEB, Cat. #M0525), and 20 units of RNase Inhibitor, Murine (NEB, Cat. #M0314). The reaction was incubated at 37 °C for 30 min, followed by heat inactivation at 80°C for 2 min. To the 15 µL reaction, 1× T4 Polynucleotide Kinase Reaction Buffer (NEB, Cat. #B0201) was added, together with 20 units of RNase Inhibitor, Murine, 55 nM [γ-³²P]-ATP (3000 Ci/mmol; Revvity, Cat. #NEG502A250UC), and 10 units of T4 PNK (NEB, Cat. #M0201). Reactions were incubated at 37 °C for 30 min and stopped by adding proteinase K (2 µg/µL in 0.5% [w/v] SDS, 10 mM EDTA pH 8.3, 20 mM Tris-HCl pH 7.5), along with 1000 cpm of an 86-mer labeled with [γ-³²P]-ATP as a recovery and loading control, and incubated at 42 °C for 15 min. Labelled RNA products were extracted with Acid-Phenol:Chloroform, pH 4.5 (with IAA, 125:24:1; Invitrogen, Cat. #AM9720), and ethanol-precipitated overnight at −20 °C. The extracted RNA was electrophoresed on a 12% (v/v) polyacrylamide (19:1) sequencing gel containing 8 M urea for 1.5 h at 80 W. Gels were dried, exposed to PhosphorImager screens, and analysed using a Typhoon FLA 9500 scanner. The western blot against FLAG tag was performed as in Western blot except a MagicMark XP Western Protein standard (Thermo Scientific, LC5602) and the secondary antibodies are horse-radish peroxidase-conjugated goat anti-mouse IgG (H + L) at 1:5000 (Thermo Scientific, 31430) were used. Secondary antibody was visualized with SuperSignal West Femto Maximum Sensitivity Substrate (Thermo Scientifi, Cat. # 34094) and scanned with ChemiDoc MP Imager (Bio-rad). Image Lab (v6.0.1) was used to quantify the intensity of both RNA sequencing gel and the western blots. The signal from the RNA was normalized by the amount of protein present in the western blot. The statistical analysis was performed as described in section Western blot.

### Molecular dynamics simulations

Atomistic molecular dynamics simulations were run with GROMACS^97^. Start structures were generated using AlphaFold2^56^ and AlphaFold3^98^ The start structures are deposited on zenodo together with the simulation parameters files (.mdp and .tpr files) and the processed molecular dynamics trajectories. Additional information about simulations is available in (Table S1). The Amber99sb-star-ildn-q protein force field^99–102^ and the TIP4P water model^103^ were used. The RNA was described with the OL3 force field^104,105^ as in the work of Collauto et al.^106^ on simulations of protein-RNA systems.

The equations of motions were integrated with a 2 fs time step. The P-LINCS algorithm^107^ was used to constrain the length of all covalent bonds. Long-range electrostatics were described by the Particle Mesh Ewald (PME) method. The grid spacing was set to 0.16 nm and a cut-off of 1 nm for real space interactions was used. To maintain the temperature at 300 K the Bussi-Dondadio-Parinello thermostat^108^ was used, with a coupling constant τ_T_ of 0.001 ps. With the Parinello-Rahman barostat^109^ we maintained the pressure at 1 bar, with a coupling constant τ_p_ =2 ps.

Energy minimisation with the steepest descent algorithm was run for 5000 steps or until the maximum force was less than 1000 kJ/mol/nm. The system was equilibrated in the NVT ensemble for 100 ps, followed by 100 ps of equilibration in the NPT ensemble. Production simulations were typically run for 1 μs or 5 μs. Thermostat and barostat settings were as in the NPT equilibration simulations.

Simulations were analysed using the MDAnalysis and mdtraj Python libraries^110–112^. Distances and contacts were computed using MDAnalysis. Secondary structure was assessed with the DSSP algorithm^113^ using mdtraj.

## DATA REPOSITORIES

All sequencing data generated for this study has been uploaded to the European Nucleotide Archive (www.ebi.ac.uk) under accession numbers PRJEB95103.

All mass spectrometry data generated in this study have been uploaded to the UCSD Mass Spectrometry Interactive Virtual Environment (MassIVE) server. In vivo N-termini profiling data are available under accession number MSV000099904 (Reviewer access ftp://MSV000099904@massive-ftp.ucsd.edu Username: MSV000099904_reviewer Password: h#BpL6$MwKKE9%ZZ). In vitro N-termini profiling data accession number MSV000099971 (Reviewer access: ftp://MSV000099971@massive-ftp.ucsd.edu Username: MSV000099971_reviewer Password: h#BpL6$MwKKE9%ZZ). DPF-3 proteomic data are available under accession number MSV000099932 (Reviewer access: ftp://MSV000099932@massive-ftp.ucsd.edu Username: MSV000099932_reviewer Password: h#BpL6$MwKKE9%ZZ). Simulation setup and simulation trajectories are available on zenodo (DOI: 10.5281/zenodo.17829502). Raw microscopy data can be found on a public Omero site: https://omero.imb.uni-mainz.de/pub/isolehto2025.

## ACKNOWLEDGEMENTS

Support by the IMB Genomics Core Facility is gratefully acknowledged. We thank the IMB Media Laboratory and Microscopy Core Facility for consumables and equipment (AF7000, DFG grant 212049334; Stellaris 8, DFG grant 497660232). We thank the microscope core facility at the Faculty of Biology of the JGU for the use of the LSM900, funded by the Ministry of Science and Health of Rhineland-Palatinate and the European Regional Development Fund (ERDF/REACT-EU, Grant No. 84012490). We thank Martin Möckel and the Protein production Core Facility at the IMB for help in making DPF-3 recombinant proteins. Florens Hillenbrand is acknowledged for his contribution in smFISH experiments. The authors gratefully acknowledge the data storage facilities provided by the Institute for Quantitative and Computational Biosciences (IQCB) at Johannes Gutenberg University Mainz. The authors gratefully acknowledge the computing time provided to them at the NHR Center NHR@SW at Johannes Gutenberg University Mainz (project nhr-dyndisphase). This is funded by the Federal Ministry of Education and Research, and the state governments participating on the basis of the resolutions of the GWK for national high performance computing at universities (www.nhr-verein.de/unsere-partner). L.S.S. acknowledges support by ReALity (Resilience, Adaptation and Longevity) and Forschungsinitiative des Landes Rheinland-Pfalz. The work was funded by the Deutsche Forschungsgemeinschaft (DFG, German Research Foundation): project numbers 464588647 (CRC1551, Polymer Concepts in Cellular Function, RFK and LS), 252386272 and 504320275 (R.F.K.).

---

**Figure S1.**
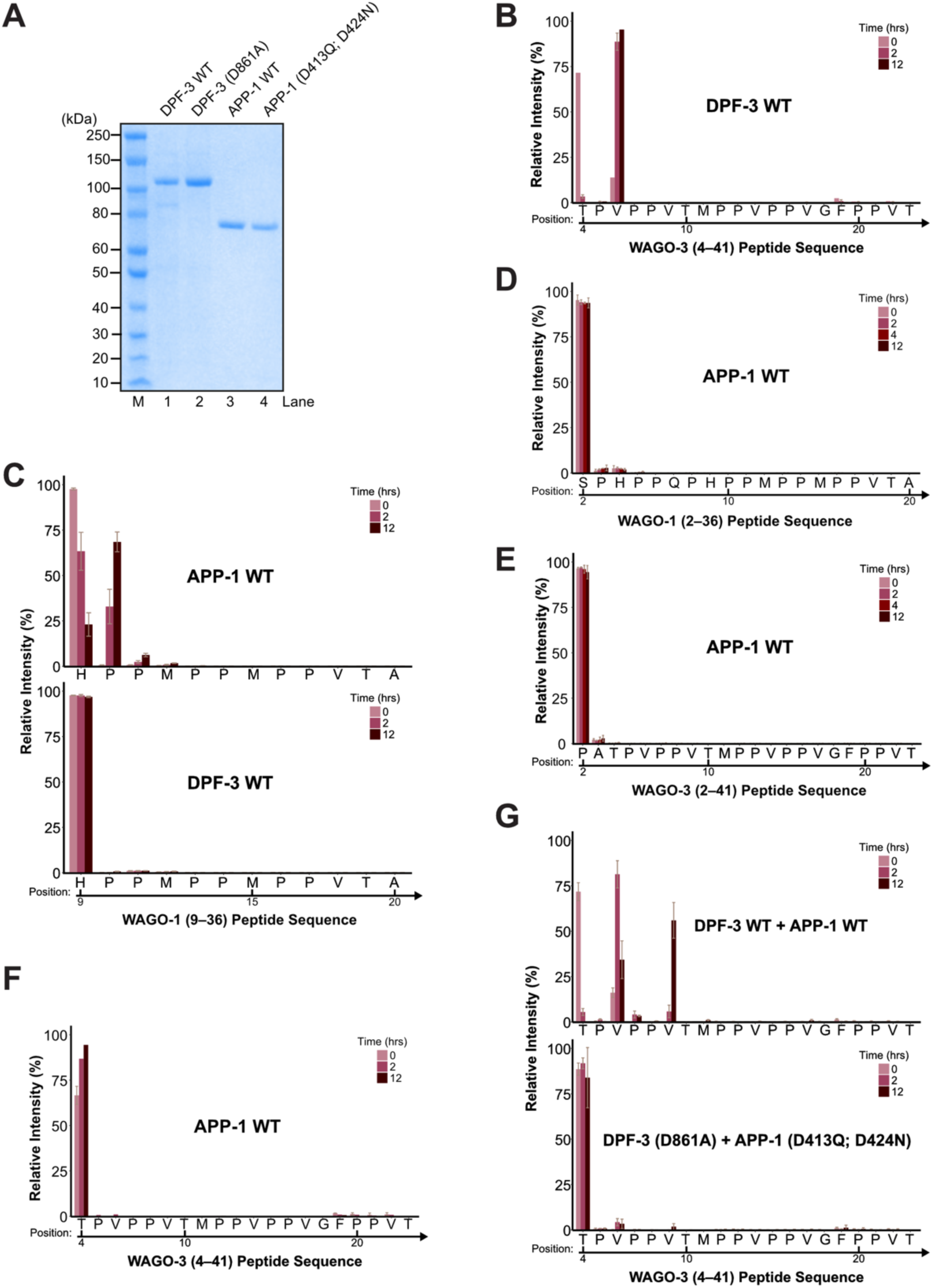
APP-1 and DPF-3 have complementary substrate preferences. (A) Coomassie-stained SDS–PAGE gel of purified recombinant DPF-3–TwinStrep and 6xHis– APP-1 proteins. The gel shows the protein stocks used for all in vitro processing assays. DPF-3 WT (lane 1), DPF-3 catalytically inactive (lane 2), APP-1 WT (lane 3), and APP-1 catalytically inactive (lane 4) each show a prominent band at the expected molecular weight. M, size marker. (B) Bar plot showing the relative peptide intensity within each sample measured by mass spectrometry. In vitro N-terminal cleavage of the WAGO-3 4aa–41aa (TPVPPVTMPPVPPVGFPPVTAPPGLHPPPPVPPVPVPTL) peptide incubated with recombinant WT DPF-3. Only a partial peptide sequence is shown on the X-axis. Numbers above the arrows indicate amino acid positions in the full-length WAGO-3 protein. The time course (0, 2, and 12 h) shows that WT DPF-3 efficiently removes the first two amino acids of WAGO-3 4aa–41aa. Bars represent means; error bars denote standard deviation (n = 2–3). The time-zero sample has only one replicate. (C) Bar plot showing the relative peptide intensity within each sample measured by mass spectrometry, as in (B). Time course of in vitro N-terminal cleavage of the WAGO-1 9aa–36aa (HPPMPPMPPVTAPPGAMTPMPPVPADAQK) peptide incubated with recombinant WT APP-1 (top) or DPF-3 (bottom). Only WT APP-1 showed consistent accumulation of one- and, to a lesser extent, two–amino acid–shorter peptides. Bars represent means; error bars denote standard deviation (n = 2–3). (D–E) Bar plots showing the relative peptide intensity within each sample measured by mass spectrometry, as in (B). Time course of in vitro N-terminal cleavage of WAGO-1 2aa–36aa (SPHPPQPHPPMPPMPPVTAPPGAMTPMPPVPADAQK) (D) and WAGO-3 2aa–41aa (PATPVPPVTMPPVPPVGFPPVTAPPGLHPPPPVPPVPVPTL) (E) peptides incubated with recombinant WT APP-1. APP-1 showed no activity, as shorter peptides were not detected for either substrate. Bars represent means; error bars denote standard deviation (n = 2–3). The time-zero sample has only one replicate. (F) Bar plot showing the relative peptide intensity within each sample measured by mass spectrometry, as in (B). Time course of *in vitro* N-terminal cleavage of the WAGO-3 4aa–41aa peptide incubated with recombinant WT APP-1. APP-1 showed no activity, as shorter peptides were not detected. Bars represent means; error bars denote standard deviation (n = 2–3). Some samples have only one replicate. (G) Bar plot showing the relative peptide intensity within each sample measured by mass spectrometry, as in (B). Time course of in vitro N-terminal cleavage of the WAGO-3 4aa–41aa peptide incubated with both recombinant WT APP-1 and DPF-3 (top) or catalytically inactive APP-1 and DPF-3 (bottom). Only the WT proteases processed several amino acids. Bars represent means; error bars denote standard deviation (n = 2–3).

**Figure S2.**
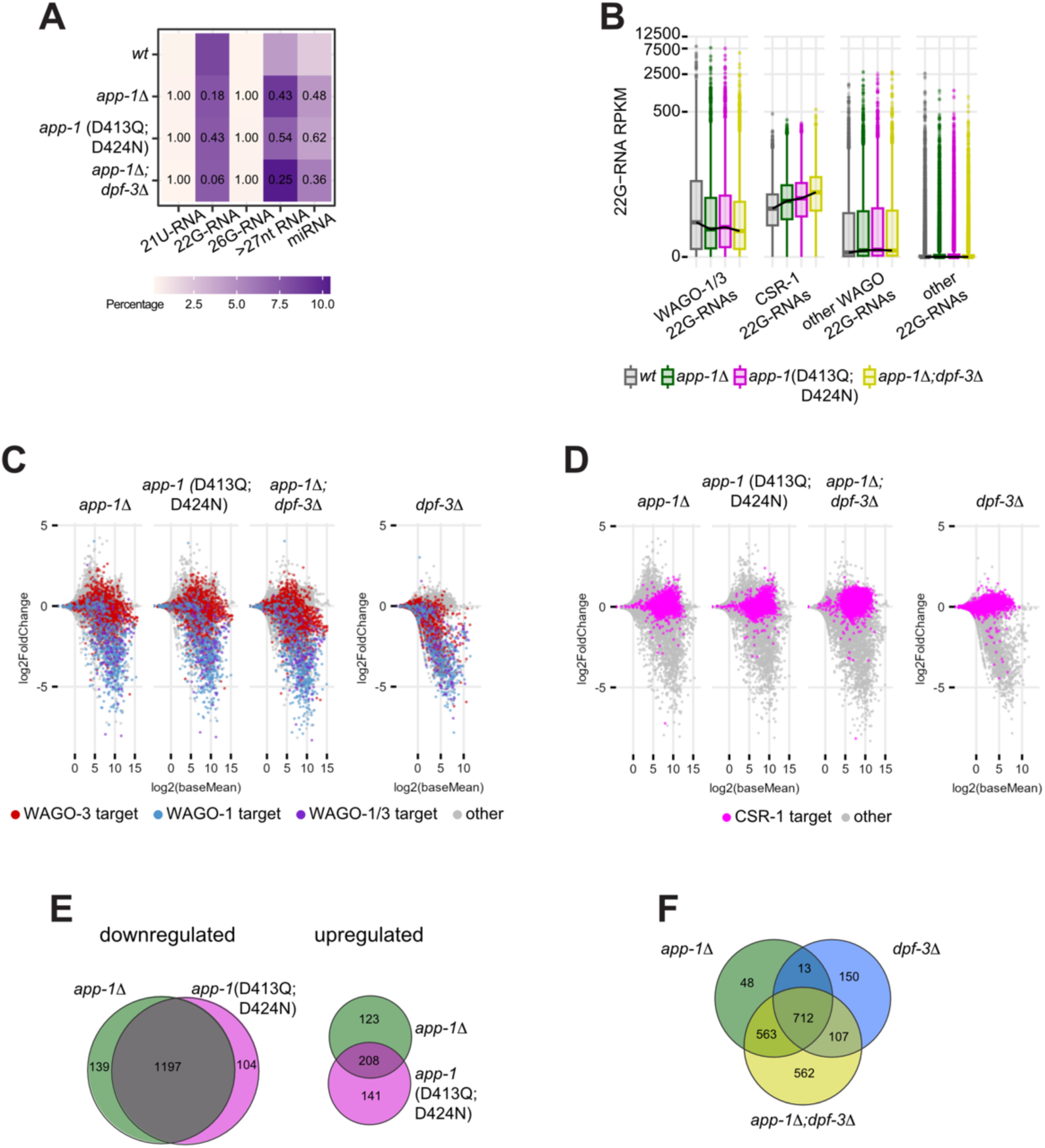
Loss of APP-1 and DPF-1 depletes WAGO-1 and WAGO-3 22G RNAs. (A) Heatmap showing relative expression of 21U-RNAs, 22G-RNAs, 26G-RNAs, miRNAs and RNAs longer than 27 nt in the indicated mutants and WT. Numbers indicate *p*-values from a Fisher#s exact test against the WT. (B) Boxplot showing the RPKMs of 22G-RNAs against WAGO-1/3 targets, CSR-1 targets, genes defined as targets of any other WAGO, and genes not previously defined as targets of any WAGO via RIP-Seq. (C) Scatterplot showing changes to 22G-RNAs in different *app-1* and *dpf-3* mutants, as indicated. *dpf-3* single mutant data from^38^. Each dot represents one gene, typically with several 22G-RNAs. Colours indicate whether each gene has been proposed as a target of WAGO-1 and/or WAGO-3 based on RIP-Seq studies^24^. (D) Scatterplot showing changes to 22G-RNAs in different *app-1* and *dpf-3* mutants, as indicated. *dpf-3* single mutant from^38^. Each dot represents one gene, typically with several 22G-RNAs. Colours indicate whether each gene has been proposed as a target of CSR-1 based on RIP-Seq studies. (E) Venn diagram showing the overlap between genes with significantly enriched 22G-RNAs and significantly decreased 22G-RNAs as compared to WT in *app-1* and *app-1*(D413Q; D424N) catalytic mutants. (F) Venn diagram of genes with significantly decreased 22G-RNA levels in *app-1*, *dpf-3*, and *app-1;dpf-3* double mutants.

**Figure S3.**
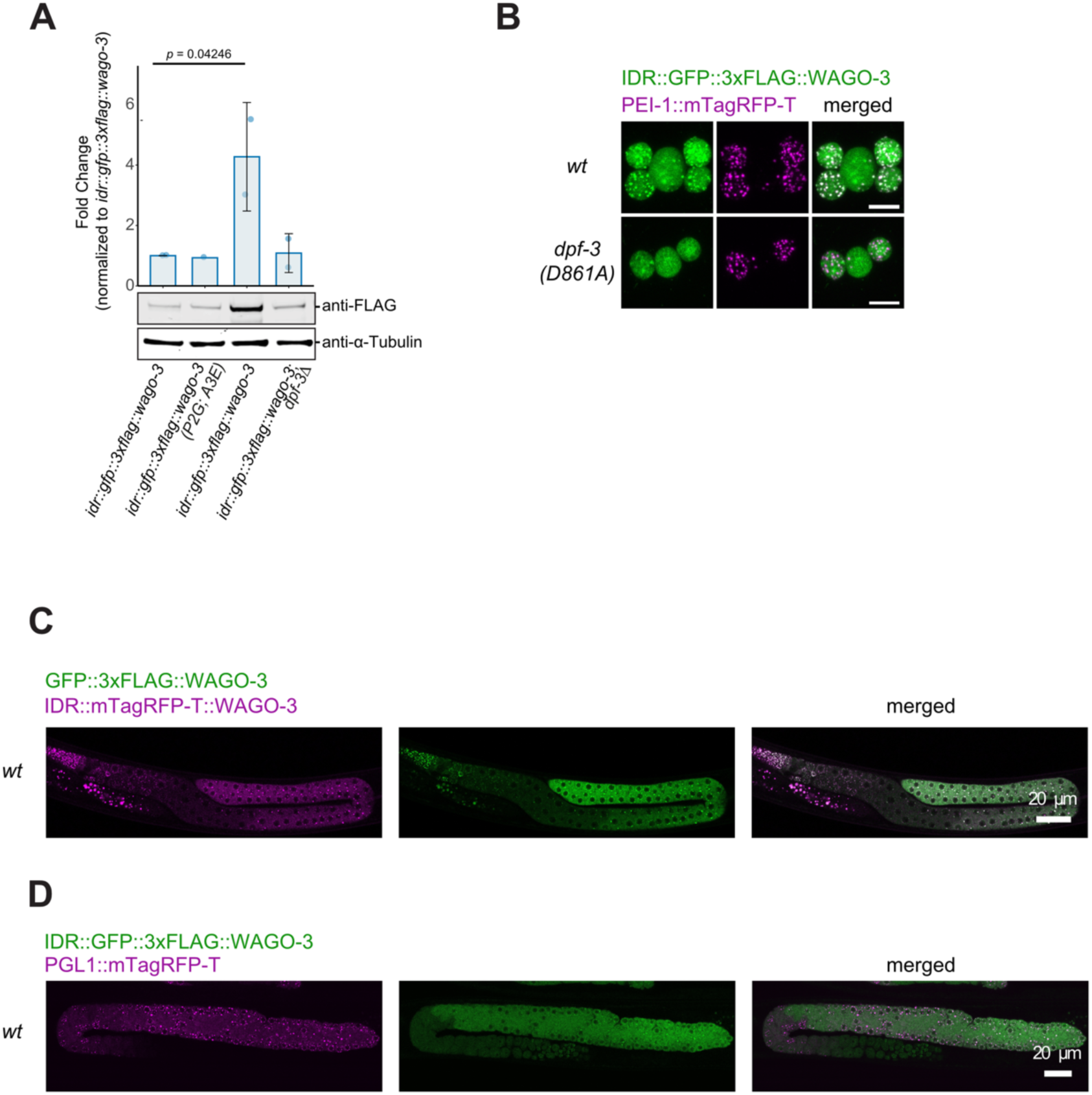
DPF-3 regulates WAGO-3 abundance and PEI granule localization independently of processing. (A) Endogenous WAGO-3 protein levels with N-terminal or internal tagging. Representative Western blot comparing levels of N-terminal 3xFLAG::GFP–tagged and internal 3xFLAG ::GFP–tagged WAGO-3 in different genetic backgrounds. Denatured whole-worm protein extracts from adult males carrying the *him-5(e1490)* mutation were resolved by SDS–PAGE. The blot (bottom) was probed with anti-FLAG to detect WAGO-3 and anti–α-tubulin as a loading control. Bar plot (top) shows fold-change values relative to the internally tagged WAGO-3. Statistical significance was determined using Welch’s two-sample t-test (p-value shown above the bars). Bars represent mean ± SD (n = 2). (B) Representative confocal images of dissected *C. elegans* budding spermatids showing IDR::GFP::3xFLAG::WAGO-3 (green), and PEI-1::mTagRFP-T (magenta) localization in the specified mutants. All strains additionally carry the *him-5(e1490)* mutation in the background. Maximum intensity projections. Scale bar, 5 μm. Images are representative of at least 10 budding spermatids pooled from at least two independent experiments. (C) Confocal image of male gonad showing N-terminally tagged GFP::3xFLAG::WAGO-3 (green), and internally tagged IDR::mTagRFP-T::WAGO-3 (magenta). The internal tagging was done at the exact same site as that for the internal GFP tag used throughout the rest of the manuscript. (D) Confocal image of L4 gonad showing internally tagged IDR::GFP::3xFLAG::WAGO-3 (green), and PGL-1::mTagRFP-T (magenta).

**Figure S4.**
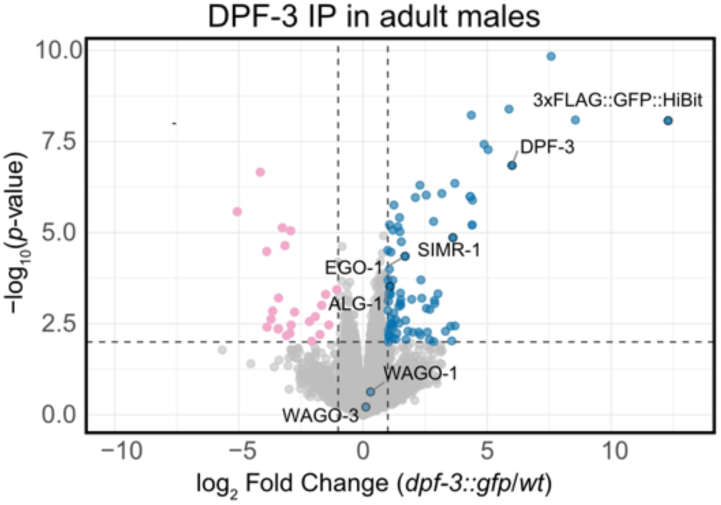
ALG-1 is enriched in DPF-3 pull-downs. Identification of DPF-3 interacting proteins. Volcano plot showing proteins enriched in DPF-3::3xFLAG::TEV::GFP::HiBit immunoprecipitations (IP), quantified using label-free proteomics. IP experiments were performed on native protein extracts from adult males (n = 4 biological replicates). The x-axis shows the mean fold change in abundance for each protein in tagged DPF-3 samples relative to untagged controls, all on a *him-5(e1490)* genetic background. The y-axis represents the −log₁₀(P) of observed enrichments. Dashed lines indicate thresholds for significance: p = 0.05 and two-fold change in enrichment. Dots indicate proteins that are enriched (light blue), depleted (pink), or detected (grey). Only uniquely matching peptides were considered. DPF-3, and known interactor are highlighted.

**Figure S5.**
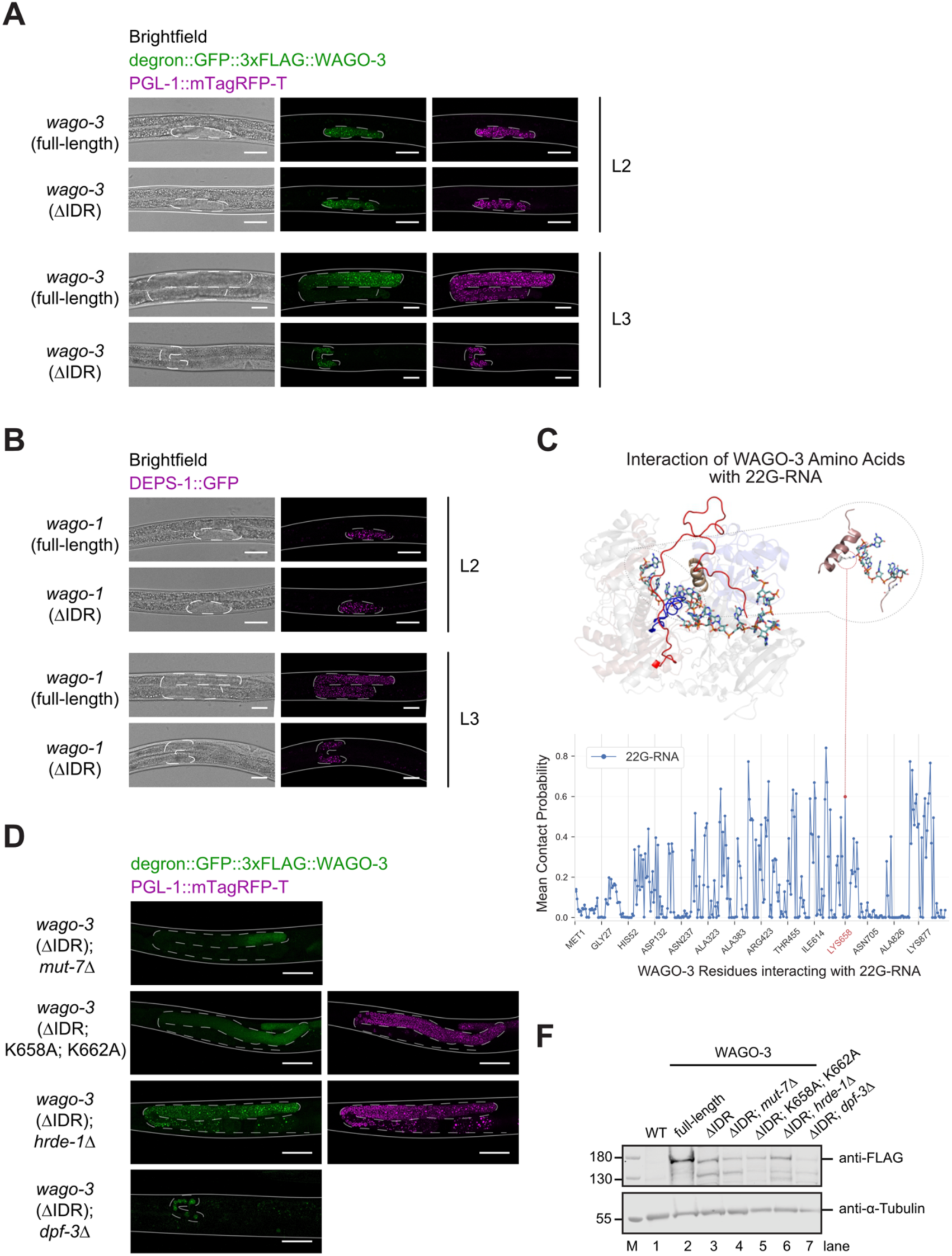
Deleting the N-IDRs from WAGO-1 and WAGO-3 leads to a sterile phenotype. (A-B) Representative fluorescence micrographs of L2- and L3-stage larvae expressing degron::GFP::3×FLAG::WAGO-3 and PGL-1::mTagRFP-T (A) and DEPS-1::GFP (B) in the indicated genetic backgrounds. Solid lines represent the worm body, and dashed lines represent the germline boundary. Images are representative of three independent biological replicates. Scale bars, 20 μm. (C) Snapshot of a wt WAGO-3 MD simulation with a 22G-RNA bound in the canonical binding cleft. The inset shows a close-up of Lys658 interaction with 5’ end of 22G-RNA. The line plot shows contact probabilities of WAGO-3 residues with 22G-RNA. (D) Representative fluorescence micrographs of day 1 adult hermaphrodites expressing degron::GFP::3×FLAG::WAGO-3 and PGL-1::mTagRFP-T in the indicated genetic backgrounds cultured on auxin-free plates. For some mutants crossing with PGL-1::mTagRFP-T led to sterility. Solid lines represent the worm body, and dashed lines represent the germline boundary. Images are representative of three independent biological replicates. Scale bars, 50 μm. (F) Immunoblot analysis of FLAG-tagged WAGO-3 levels in the indicated genetic backgrounds cultured on auxin-free plates. Whole-worm protein extracts from gravid adult hermaphrodites were resolved by SDS-PAGE and probed with anti-FLAG antibody. α-tubulin served as a loading control. n = 1.

**Figure S6.**
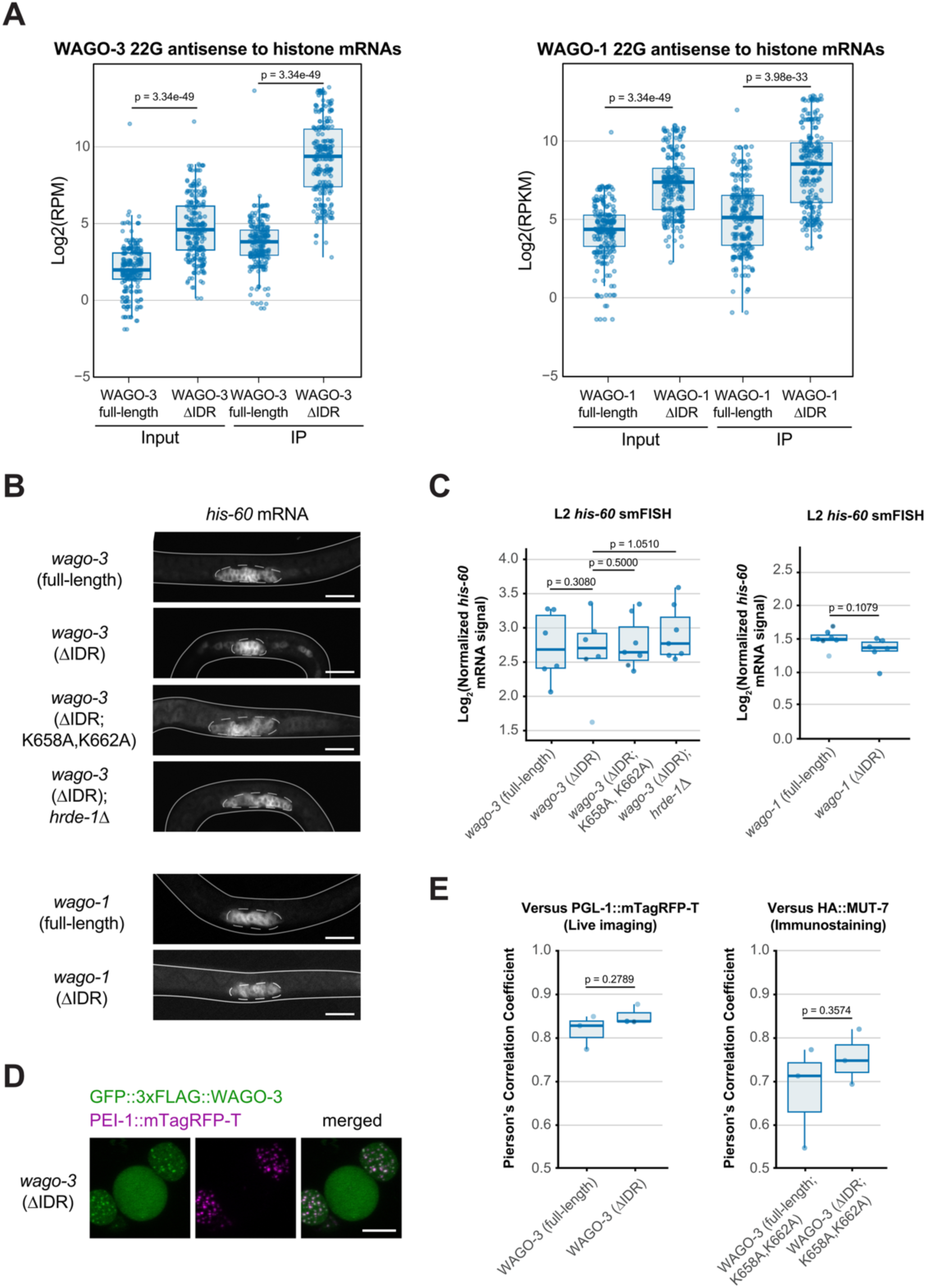
The binding of 22G-RNAs antisense to histone mRNAs by WAGO-1/3 (ΔIDR) has no effect in L2 stage worms. (A) Abundance of histone gene-targeting 22G-RNAs in input and immunoprecipitated samples. Log₂-transformed reads per million (RPM) values for 22G-RNAs mapping to histone genes are shown. Each dot represents an individual histone gene; horizontal bars indicate mean values, and error bars represent standard deviation. Statistical significance was determined using an unpaired two-sample Wilcoxon test. n = 4 biological replicates per genotype. (B) Maximum intensity projections of confocal *his-60* smFISH in L2-stage larvae cultured on auxin-free plates. Lines outline the animal and dashed lines outline the germline. Images are representative of six biological replicates. Scale bar, 20 μm. (C) Quantification of normalized *his-60* mRNA smFISH signals in L2-stage larvae shown in (B). Each dot represents an individual animal; horizontal bars indicate mean values, and error bars represent standard deviation. Statistical significance was assessed using Welch’s t-test for WAGO-1 comparisons and Dunnett’s multiple comparison test for WAGO-3 comparisons. n = 6 biological replicates per genotype. (D) Representative confocal images of dissected *C. elegans* budding spermatids showing GFP::3xFLAG::WAGO-3 (green), and PEI-1::mTagRFP-T (magenta) localization in the specified mutants. All strains additionally carry the *him-5(e1490)* mutation in the background. Maximum intensity projections. Scale bar, 5 μm. Images are representative of at least 10 budding spermatids pooled from at least two independent experiments. (E) Quantification of protein colocalization with P granule and Mutator foci components. Pearson’s correlation coefficient values were calculated to assess colocalization between the indicated proteins and PGL-1 or MUT-7, respectively. Measurements were obtained from perinuclear granules across three animals per condition. Each dot represents an individual animal; horizontal bars indicate mean values, and error bars represent standard deviation. Statistical significance was determined using Welch’s t-test. n = 3 biological replicates per genotype.

**Figure S7.**
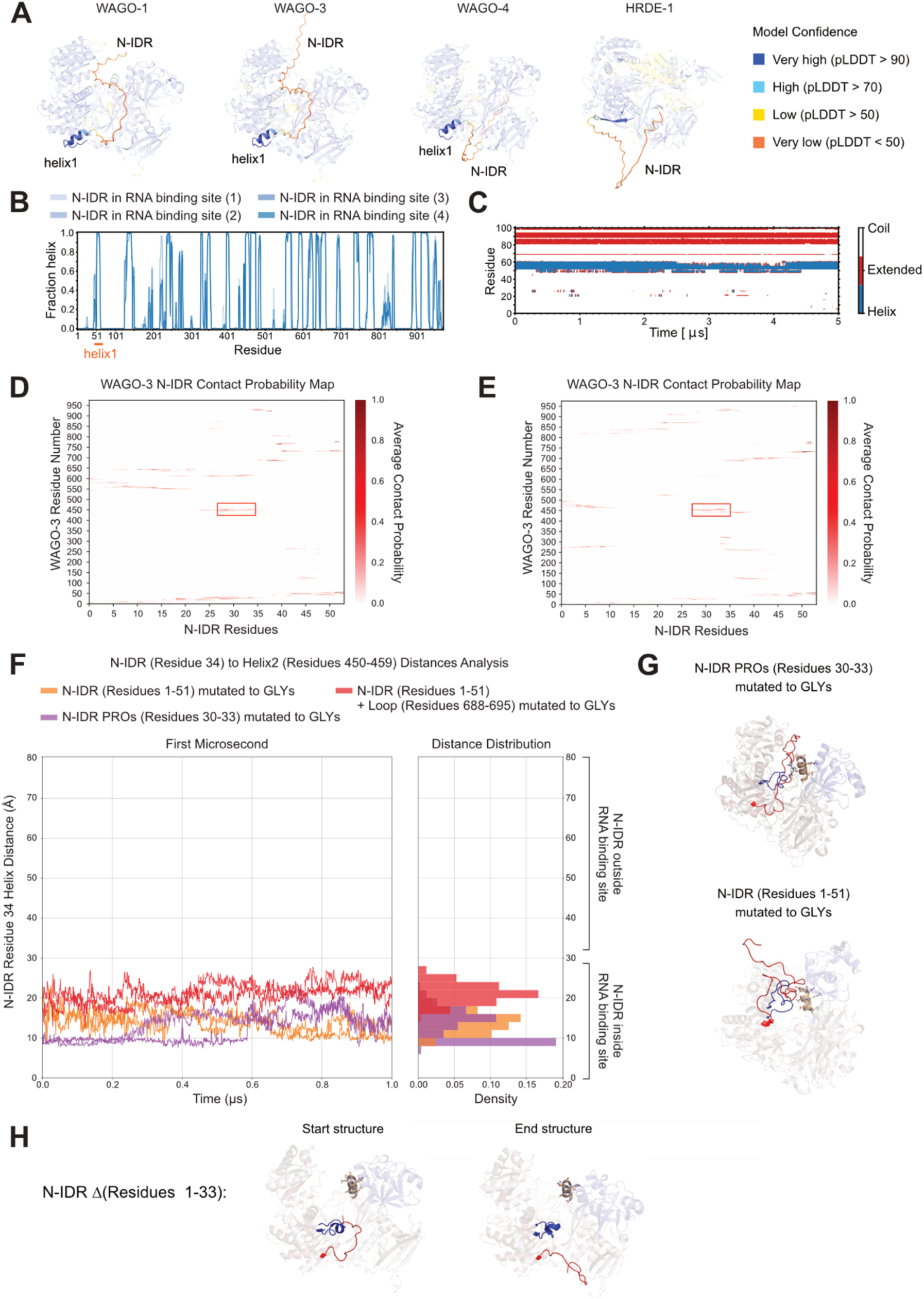
Molecular dynamics simulations show the N-IDR positioned within the RNA-binding groove. (A) AlphaFold2 structure predictions for worm-specific Argonautes representing the N-IDR and alpha-helix^1^ at the end of the N-IDR. Colors represent model confidence. (B) Fraction of time spent in helical conformations. DSSP secondary structure assignments per amino acid residue were averaged over the 5 µs simulation duration. Four trajectories, started from two different AlphaFold model are shown. (C) Secondary structure of WAGO-3 residues 1-100 assessed over the course of a 5 µs molecular dynamics simulation trajectory. DSSP was used to assign the secondary structure of individual residues. (D-E) Two replicates of contact probability maps of wt WAGO-3 simulations showing residue-level contact probabilities between WAGO-3 and the IDR (heatmap color coded from 0 (no contact) to 1.0 (persistent contact)). (F) Multi-group comparison of distances between N-IDR residue 34 and helix^2^ residues 450-459 across simulations lasting 1 μs. Different simulation conditions are color-coded, individual replicates are indicated by transparency. (G) Representative structures from a 1-μs simulation of N-IDR mutations. The N-IDR is shown in red, with helix^1^ highlighted in blue, the interacting helix^2^ (residues 450-459) is represented in wheat, loop (residues 688-695) is in green. (H) Representative start and end structures from a 1-μs simulation of a truncated N-IDR variant lacking the proline residues. Removal of the proline cluster destabilizes the N-IDR– helix^2^ interaction and alters N-IDR positioning over the course of the simulation. The N-IDR is shown in red, with helix^1^ highlighted in blue, the interacting helix^2^ (residues 450-459) is represented in wheat, loop (residues 688-695) is in green.

**Table S1.** Atomistic molecular simulations generated in this study. Overview of molecular dynamics simulation setups. Simulations characteristics, simulation model (force field), box geometry, distance to periodic box edges, system charge, and simulation durations are listed. Simulations that can be retrieved from zenodo (DOI: 10.5281/zenodo.17829502).

| Supplementary Simulation | force field | Water model | Box type | Min protein to box-edge distance | System charge | time (μs) |
| --- | --- | --- | --- | --- | --- | --- |
| N-IDR inside RNA-binding site 1 | AMBER99SB-star-ildn-q | TIP4P | dodecahedron | 1.5 nm | neutral | 5 |
| N-IDR inside RNA-binding site 2 | AMBER99SB-star-ildn-q | TIP4P | dodecahedron | 1.5 nm | neutral | 5 |
| N-IDR inside RNA-binding site 3 | AMBER99SB-star-ildn-q | TIP4P | dodecahedron | 1.5 nm | neutral | 5 |
| N-IDR inside RNA-binding site 4 | AMBER99SB-star-ildn-q | TIP4P | dodecahedron | 1.5 nm | neutral | 5 |
| N-IDR outside RNA-binding site 1 | AMBER99SB-star-ildn-q | TIP4P | dodecahedron | 1.5 nm | neutral | 5 |
| N-IDR outside RNA-binding site 2 | AMBER99SB-star-ildn-q | TIP4P | dodecahedron | 1.5 nm | neutral | 5 |
| 22G-RNA in RNA-binding site 1 | AMBER99SB-star-ildn-q | TIP4P | dodecahedron | 1.5 nm | neutral | 2 |
| 22G-RNA in RNA-binding site 2 | AMBER99SB-star-ildn-q | TIP4P | dodecahedron | 1.5 nm | neutral | 2 |
| 22G-RNA in RNA-binding site 3 | AMBER99SB-star-ildn-q | TIP4P | dodecahedron | 1.5 nm | neutral | 2 |
| 22G-RNA in RNA-binding site 4 | AMBER99SB-star-ildn-q | TIP4P | dodecahedron | 1.5 nm | neutral | 2 |
| 22G-RNA in RNA-binding site 5 | AMBER99SB-star-ildn-q | TIP4P | dodecahedron | 1.5 nm | neutral | 2 |
| 22G-RNA in RNA-binding site 6 | AMBER99SB-star-ildn-q | TIP4P | dodecahedron | 1.5 nm | neutral | 2 |
| 22G-RNA in RNA-binding site 7 | AMBER99SB-star-ildn-q | TIP4P | dodecahedron | 1.5 nm | neutral | 2 |
| 22G-RNA in RNA-binding site 8 | AMBER99SB-star-ildn-q | TIP4P | dodecahedron | 1.5 nm | neutral | 2 |
| N-IDR Δ(Residues 1-29) 1 | AMBER99SB-star-ildn-q | TIP4P | dodecahedron | 1.5 nm | neutral | 1 |
| N-IDR Δ(Residues 1-29) 2 | AMBER99SB-star-ildn-q | TIP4P | dodecahedron | 1.5 nm | neutral | 1 |
| N-IDR Δ(Residues 1-33) 1 | AMBER99SB-star-ildn-q | TIP4P | dodecahedron | 1.5 nm | neutral | 1 |
| N-IDR Δ(Residues 1-33) 2 | AMBER99SB-star-ildn-q | TIP4P | dodecahedron | 1.5 nm | neutral | 1 |
| N-IDR PRO (Residues 30-33) mutated to GLY 1 | AMBER99SB-star-ildn-q | TIP4P | dodecahedron | 1.5 nm | neutral | 1 |
| N-IDR PRO (Residues 30-33) mutated to GLY 2 | AMBER99SB-star-ildn-q | TIP4P | dodecahedron | 1.5 nm | neutral | 1 |
| N-IDR (Residues 1-51) mutated to GLY 1 | AMBER99SB-star-ildn-q | TIP4P | dodecahedron | 1.5 nm | neutral | 1 |
| N-IDR (Residues 1-51) mutated to GLY 2 | AMBER99SB-star-ildn-q | TIP4P | dodecahedron | 1.5 nm | neutral | 1 |
| N-IDR (Residues 1-51) + Loop (Residues 688-695) mutated to GLY 1 | AMBER99SB-star-ildn-q | TIP4P | dodecahedron | 1.5 nm | neutral | 1 |
| N-IDR (Residues 1-51) + Loop (Residues 688-695) mutated to GLY 2 | AMBER99SB-star-ildn-q | TIP4P | dodecahedron | 1.5 nm | neutral | 1 |

## Movie S1. WAGO-3 dynamics

This movie displays the dynamics of WAGO-3, as determined by atomistic molecular dynamics simulations. In the movie different elements are colour-coded: Blue: Helix^1^; Red: N-IDR; wheat: Helix^2^; olive: Loop (688-695).

## Movie S2. WAGO-3 dynamics with mutant N-IDR

This movie displays the dynamics of WAGO-3 in which Pro 30-33 have been mutated to glycine, as determined by atomistic molecular dynamics simulations. In the movie different elements are colour-coded: Blue: Helix^1^; Red: N-IDR; Wheat: Helix^2^; Olive: Loop (688-695).

